# Nature AND Nurture: Enabling formate-dependent growth in *Methanosarcina acetivorans*

**DOI:** 10.1101/2024.01.08.574737

**Authors:** Jichen Bao, Tejas Somvanshi, Yufang Tian, Maxime Laird, Pierre Simon Garcia, Christian Schöne, Michael Rother, Guillaume Borrel, Silvan Scheller

## Abstract

Methanogenic archaea are crucial in global carbon cycling as around 1 Gt of the potent greenhouse gas, methane, is produced annually. Major contributors belong to the order Methanosarcinales, which contain some of the most versatile methanogens that are capable of acetotrophic, methylotrophic and CO_2_-reducing methanogenesis. The genetically tractable model methanogen, *Methanosarcina acetivorans*, by its nature shows versatility in substrate utilization and energy conservation pathways but cannot utilize formate. In this study, we expanded the primary metabolism of *M. acetivorans* to include formate-dependent methanogenesis. By introducing an exogenous formate dehydrogenase, the two metabolically engineered *M. acetivorans* strains acquired the capacity for formate-dependent methanogenesis pathways with one capable of formate-dependent methyl-reduction and the other capable of formate-dependent CO_2_-reduction. Through nurturing the strain capable of CO_2_-reduction with adaptive laboratory evolution, we were able to enable growth and methanogenesis of *M. acetivorans* solely on formate, a metabolism only reported in methanogens without cytochromes which are limited by their versatility. *M. acetivorans* also showed acetogenic potential where the formate-dependent CO_2_-reducing strain was able to divert ≈ 10% of carbon to acetate instead of methane. Our results show that even though *M. acetivorans* lacks energy converting hydrogenase and cannot use H_2_, it has yet-uncharacterized capacity to obtain reduced ferredoxins from oxidizing formate. Our work encourages reevaluation of our understanding of formate utilization in Methanosarcinales. By enabling formate-dependent methanogenesis, we have expanded the substrate spectrum of a versatile model methanogen with cytochromes to include formate as well.

## Introduction

Methanogenic archaea play a significant role in the global carbon cycle, responsible for around 1 gigaton of methane production annually(1).

Methanogenesis usually proceeds with a substrate being reduced to methane using electron donor in an energy conserving manner that yields ATP. Along with ATP, reduced ferredoxins (Fd_red_) are required for anabolic reactions and in the first step in CO_2_-reducing methanogenesis i.e., CO_2_ reduction on methanofuran (See examples in Fig. 1, Fig. S1) (2–4). The electron donor is typically H_2_ or the carbon source itself with exceptions for secondary alcohols (2, 5). The last step for methane production is common and catalyzed by methyl-coenzyme M reductase (Mcr), which generates the heterodisulfide (HDS) of coenzyme M (CoM) and coenzyme B (CoB) as a second product that is crucial for energy conservation. This low potential HDS is reduced by heterodisulfide reductase (Hdr) in an energy conserving manner by either flavin-based electron bifurcation via HdrABC or cytochrome-dependent electron transfer via HdrED. The other energy conserving step during methanogenesis occurs while transferring the methyl group from tetrahydrosarcinapterin (H_4_SPT) to the thiol CoM, catalyzed by methyl-H_4_SPT:HS-CoM methyltransferase (Mtr), by generating a sodium motive force. The proton and sodium motive force, generated indirectly by flavin-based electron bifurcation or directly by cytochrome-dependent electron transfer along with Mtr, is used for synthesis of ATP and Fd_red_ synthesis in methanogenesis (3).

**Figure 1.**
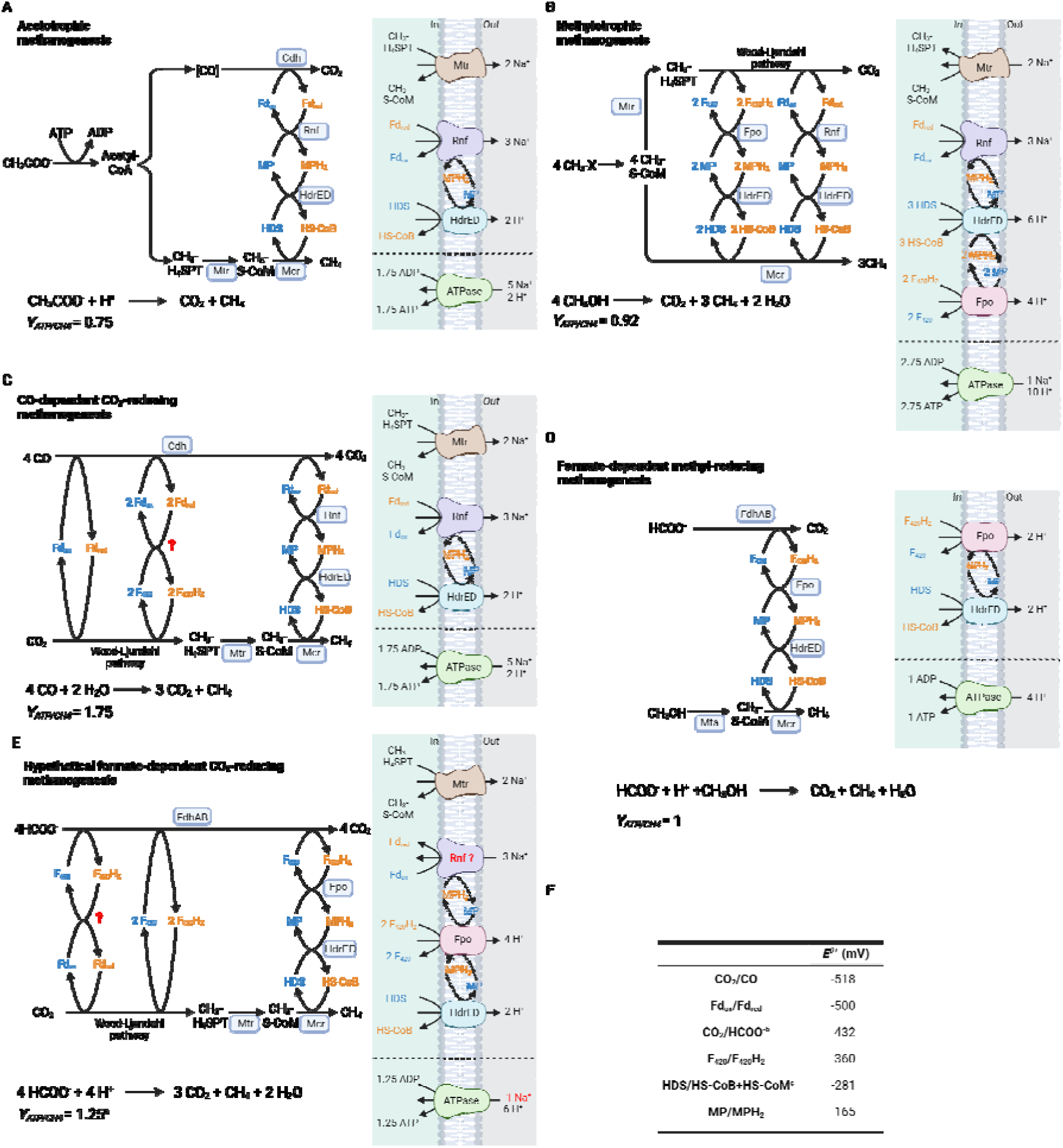
The three major methanogenesis pathways (A-C) in *M. acetivorans* and the two newly engineered pathways (D-E) (A) Acetotrophic methanogenesis - the carboxyl carbon of acetate is oxidized to obtain reduced ferredoxin. This reduced ferredoxin is then used to reduce heterodisulfide. (B) Methylotrophic methanogenesis - 1 molecule of methanol is oxidized to CO_2_ to provide 2 F_420_H_2_ and 1 Fd_red_ that are then used to reduce 3 HDS formed from 3 methanol reduced to methane. (C) CO-dependent CO_2_-reducing methanogenesis, Fd_red_ are obtained from oxidizing CO to CO_2_. Unlike *M. barkeri*, where H_2_ can be used as an intermediate, *M. acetivorans* cannot use H_2_ (D) Engineered formate-dependent methyl-reducing methanogenesis-formate is oxidized to reduce the HDS formed from methanol reduced to methane. (E) Engineered formate-dependent CO_2_-reducing methanogenesis-formate disproportionation leads to formation of 3 CO_2_ and 1 CH_4_. The yet-unknown step (marked in red question mark) is how Fd_red_ is obtained from formate oxidation for CO_2_ reduction. One of the probable hypotheses with Rnf shown. (F) Standard reduction potentials of involved redox cofactors. All reduced cofactors are in orange and the oxidized counterparts are in blue. Chemical equations and ATP yield per methane produced from respective pathways at the bottom. For simplicity, some reactants (e.g. free HS-CoM) not shown. ^a^ATP yield as calculated from the hypothetical Rnf-based energy pathway (Details in SI Appendix text and Fig S8). ^b^E’ for CO_2_/HCOO^-^ can be much higher in physiological conditions ^c^E^’m^ for HDS = -192 mV (4). Can also be higher based on actual in vivo concentrations of HDS/HS-CoM+HS-CoB. Abbreviations-Coenzyme M (CoM), Coenzyme B (CoB), heterodisulfide between CoM and CoB (HDS), methanophenazine (MP), ferredoxin (Fd), carbon monoxide dehydrogenase (Cdh), methyl-H_4_SPT:HS-CoM methyltransferase (Mtr), F_420_H_2_ dehydrogenase (Fpo), heterodisulfide reductase (Hdr), *Rhodobacter* nitrogen fixation complex (Rnf), formate dehydrogenase (Fdh).

*Methanosarcina acetivorans* is one of the most versatile methanogens in its substrate utilization (methanol, methyl amines, methyl sulfides, carbon monoxide, acetate) (6–9). It is also genetically tractable, with various genetic tools available, making it a model organism for future applications in biofuel production and even methane oxidation (10–13). Versatility of *M. acetivorans* spans all major methanogenesis pathways summarized in Fig. 1 - (A) acetotrophic, (B) methylotrophic and (C) CO-dependent CO_2_-reducing methanogenesis.

Formate can replace H_2_ in methanogenesis using formate dehydrogenase. This has been shown or proposed in various archaeal clades such Methanococcales, Methanomicrobiales or Methanonatronarchaeales (9, 14). Formate is an attractive energy carrier given that it is much easier to handle, transport, and store compared to hydrogen gas (15). Formate along with other one-carbon compounds such methanol can be derived from CO_2_ and renewable energy with high efficiency. Biological platforms capable of integrating methanol and formate efficiently into their metabolism could prove effective in CO_2_ valorization into biofuels and other value-added products (16). Intriguingly, Methanosarcinales have never been shown to use formate, although some strains of *Methanosarcina barkeri* and *Methanosarcina mazei* have genes annotated as F_420_-reducing formate dehydrogenase (FdhAB) (17, 18). FdhB reduces F_420_ using electrons from oxidation of formate by FdhA (18). While its formate oxidation activity was demonstrated, *M. barkeri* Fusaro was incapable of growth on formate as sole carbon source (18, 19). The role of FdhAB in Methanosarcinales thus remains elusive and its use could provide an additional microbe as a tool in biotechnology that can efficiently incorporate formate and methanol in its central metabolism.

When growing on formate, the microbe would have access to F_420_H_2_ but would have to synthesize Fd_red_. Synthesis of Fd_red_ (*E*°’ = -500 mV) from F_420_H_2_ (*E*°’ = -360 mV) is endergonic (Fig. 1F) and requires coupling to an exergonic reaction or energy input in terms of ion gradient (1). In methanogens without cytochromes flavin-based electron bifurcation along with energy-converting hydrogenase (Eha/Ehb) enables availability of Fd_red_ for CO_2_ reduction and anabolic reactions (20). In methanogens with cytochromes, as shown in *M. barkeri*, a different energy-converting hydrogenase (Ech) is essential to make Fd_red_ available for CO_2_ reduction and for anabolic reactions (SI Appendix Fig. S1) (20). In both cases, H_2_ is used as an intermediate for the generation of Fd_red_. *M. acetivorans* lacks the amenity of using H_2_ as intermediate due to the lack of Ech hydrogenase and hence would have to employ a yet unknown strategy to survive on formate (21).

We explored the potential use of formate dehydrogenase in *M. acetivorans* in two new strains heterologously expressing FdhAB from *M. barkeri. M. acetivorans* JBA01 (*mtr::fdhAB*) was able to grow in methanol and formate (with and without added acetate and pyruvate), reflecting its ability to use formate as electron donor. *M. acetivorans* JBAF02 (*frh::fdhAB*) was able to grow solely on formate, a metabolism that was previously reported only for methanogens without cytochromes. This study showcases CO_2_ reduction in *M. acetivorans* that is possible without providing CO. Both strains (JBA01 and JBAF02) showcase the redundancy in the energy metabolism of *M. acetivorans* to generate Fd_red_. We propose different possibilities for energy conservation in the microbe when growing on formate (Fig. 1D-E). Phylogenetic analysis indicates that Methanosarcinales acquired FdhA several times independently, probably by horizontal gene transfers (HGTs). Our results highlight the remarkable metabolic flexibility of Methanosarcinales, both in an artificial and natural way. This work enriches the panel of metabolic abilities of *Methanosarcina* which can find several applications in biofuel/value-added product production and global efforts to sustainably balance the carbon cycle to address climate change.

## Results

### Distribution and phylogeny of the F_420_-reducing formate dehydrogenase

To enable formate-dependent methanogenesis in *M. acetivorans* WWM73, a well characterized FdhAB was needed for heterologous expression.

Homologues of FdhA were searched in a database comprising 9,596 bacterial and 1,268 archaeal proteomes, including 68 Methanosarcinales. FdhA is highly prevalent among methanogens belonging to the lineages most closely related to the Methanosarcinales, *i*.*e*., Methanocellales, Methanoflorentales (Bog-38), Methanomicrobiales and Halobacteriales (Fig. 2A). In contrast, Methanosarcinales exhibit an uneven distribution of FdhA, with a comparatively lower number of genomes encoding it. On one hand, all Methanotrichaceae (a basal lineage in Methanosarcinales) genomes contain at least one *fdhA* gene and most of them have between three and seven copies of this gene. On the other hand, FdhA is only present in 8 out of the 43 Methanosarcinaceae in our database (Fig. 2A). The phylogeny of FdhA revealed that this gene was acquired multiple times through HGT during the evolution of Methanosarcinales (Fig. 2B). These HGT in Methanosarcinaceae are further supported by the patchy distribution of the enzyme in this family. Indeed, vertical inheritance in Methanosarcinaceae species is unlikely as it would imply a large number of losses, including many recent losses in *Methanosarcina*. The phylogeny of FdhA also indicates that the multiple copies present in Methanotrichaceae originate from several HGTs rather than gene duplication (Fig. 2B).

**Figure 2.**
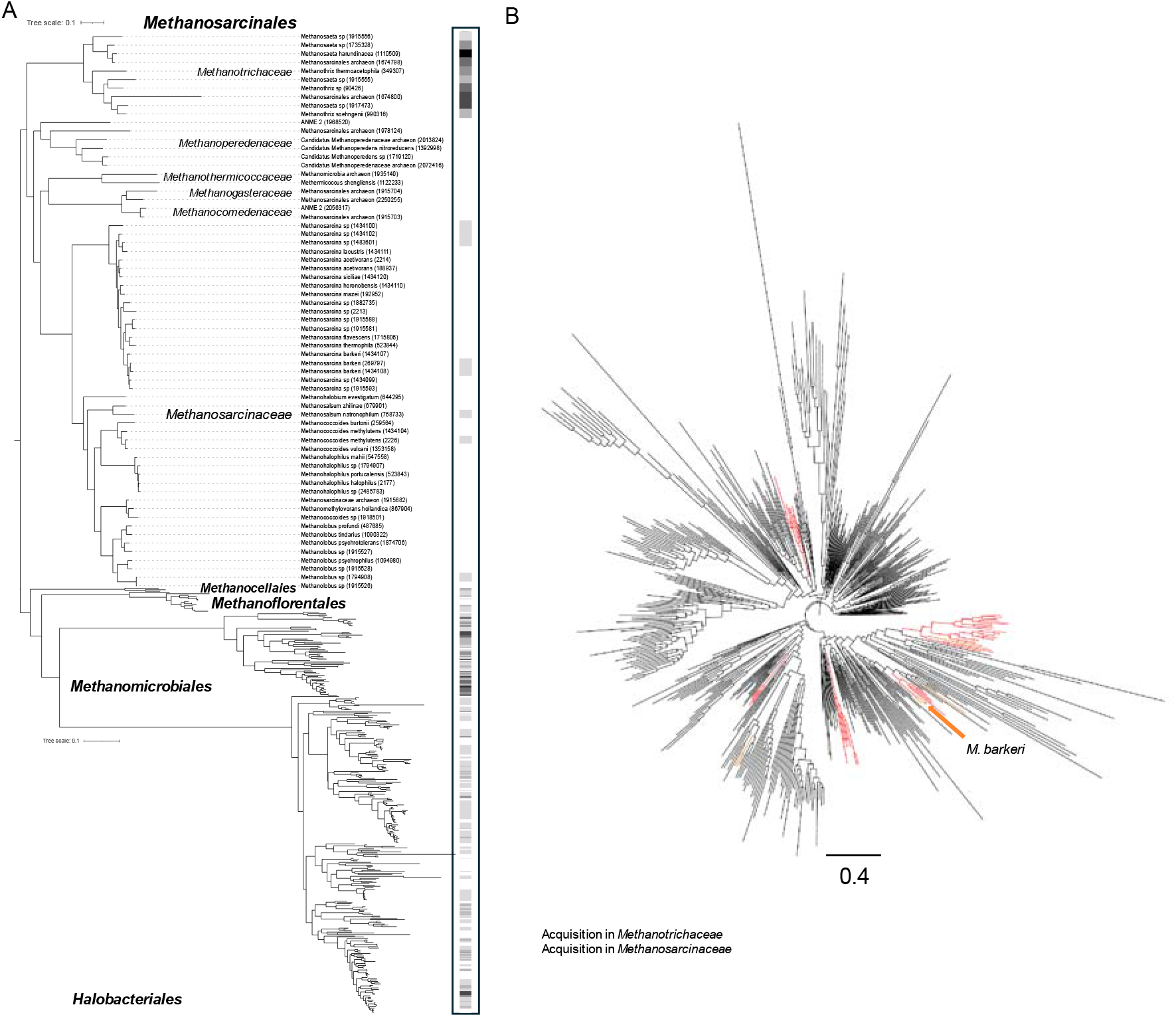
Taxonomic distribution and phylogenetic tree FdhA. A. Taxonomic distribution of FdhA in class II methanogens and *Halobacteriales*. The phylogenetic tree has been inferred based on a concatenation of If2, RpoB and RpoC alignments (LG+F+R10+C20+PMSF, 383 sequences, 2,523 amino-acid positions). The dots at the nodes correspond to ultrafast-bootstrap values > 95%. A zoom on the tree has been applied for Methanosarcinales. The number of copies of FdhA is represented by a gray gradient (white = 0, black = 7). The numbers in parenthesis at the tips correspond to the NCBI taxonomic ID. The scale bars represent the average number of substitutions per site. B. Phylogenetic tree of FdhA (LG+R10, 540 sequences, 658 amino-acid positions). Stars represent independent acquisitions in Methanosarcinales. A subsampling of sequences has been applied for non-Methanosarcinales sequences. The scale bars represent the average number of substitutions per site.

FdhAB from *M. barkeri* Fusaro was chosen for heterologous expression as *M. barkeri* is closely related to *M. acetivorans* and, unlike FdhAB from *Methanococcus maripaludis*, the FdhAB from *M. barkeri* does not contain selenocysteine. Growth of *M. barkeri* on methanol and formate together showed higher methane levels in the presence of methanol and formate than with methanol alone where formate utilization was observed through NMR quantification (SI Appendix Table S1) confirming an active FdhAB and formate utilization for methanogenesis in *M. barkeri*.

For further characterization, the enzyme was heterologously produced in *M. acetivorans* WWM73. *fdhAB* was expressed from a plasmid under the control of P*mcrB*(tetO1) promoter (22). The activity of the recombinant enzyme was assessed by measuring its ability to reduce benzyl viologen in cell-free extracts. The enzyme exhibited an activity of 0.72 U mg^-1^ of total protein (SI Appendix Fig. S2), confirming functional expression in *M. acetivorans*.

### Effect of formate on *M*. acetivorans

To gather physiological reference before incorporating formate dehydrogenase, *M. acetivorans* WWM73 was cultivated with and without added formate with methanol as the carbon source. The strain did not consume any formate (SI Appendix Fig. S3D) and the final methane yield from methanol remained unchanged regardless of whether formate was present or not (SI Appendix Fig. S3B-C). *M. acetivorans* grew slightly slower in the presence of formate (doubling time ≈ 8 hours) than in its absence (doubling time ≈ 7 hours) (SI Appendix Fig. S3A).

Differential gene expression analysis of *M. acetivorans* WWM73 grown with and without formate revealed 836 differentially expressed genes (DEGs) (p_adj_ < 0.01) that exhibited significant upregulation or downregulation (|Log2FC| > 0.5) (SI Appendix Table S2 and Dataset S1) out of a total of 4544 coding sequences. 205/836 genes were annotated as hypothetical proteins, suggesting their potential involvement in yet-uncharacterized processes. 11 genes were classified as transcriptional regulators along with 9 sensory transduction histidine kinases.

*ma0220* (*fdhD*, gene encoding sulfur carrier protein required for Fdh activity), although differentially expressed (p_adj_ < 0.01), was not significantly different (Log2FC = -0.25) in presence of formate. *ma4008, ma0103* and *ma4393* are the members of the Acetate uptake Transporter (AceTr) family, which has been reported to transport carboxylic acids. Only *ma4008* (*aceP*, characterized acetate specific transporter (23)) was upregulated, to support acetate efflux, as higher acetate production (also reflected in upregulated *cdh, ack* and *pta)* was seen in presence of formate (SI Appendix Fig. S3E). *ma0103* and *ma4393* were not differentially expressed. *fpo, hdrED, hdrC2* were also upregulated in presence of formate (SI Appendix Fig. S3F).

### Formate-dependent methyl-reducing methanogenesis

*M. acetivorans* JBA01 was constructed by inserting *fdhAB* from *M. barkeri* into the *mtr* operon, utilizing the promoter of *mtr* for transcription and additional terminators at the end ensuring that *mtr* is disrupted and not expressed. Without Mtr, the strain lost its ability to grow solely on methanol but could grow mixotrophically on methanol and acetate, like another previously characterized *M. acetivorans mtr* mutant (24). However, with functional FdhAB, the strain could also utilize formate instead of acetate.

The doubling time of *M. acetivorans* in MFAcP media (60 mM methanol and formate; 5 mM acetate and pyruvate) was ≈ 11 hours (Fig. 3A). Acetate and pyruvate were added to the media for anabolic reactions as Fd_red_ could not be obtained via methanol oxidation in absence of Mtr. Metabolite analysis showed that all formate, methanol, and acetate were consumed by the end of growth, and the methane yield reached the theoretical value. (Fig. 3B-E).

**Figure 3.**
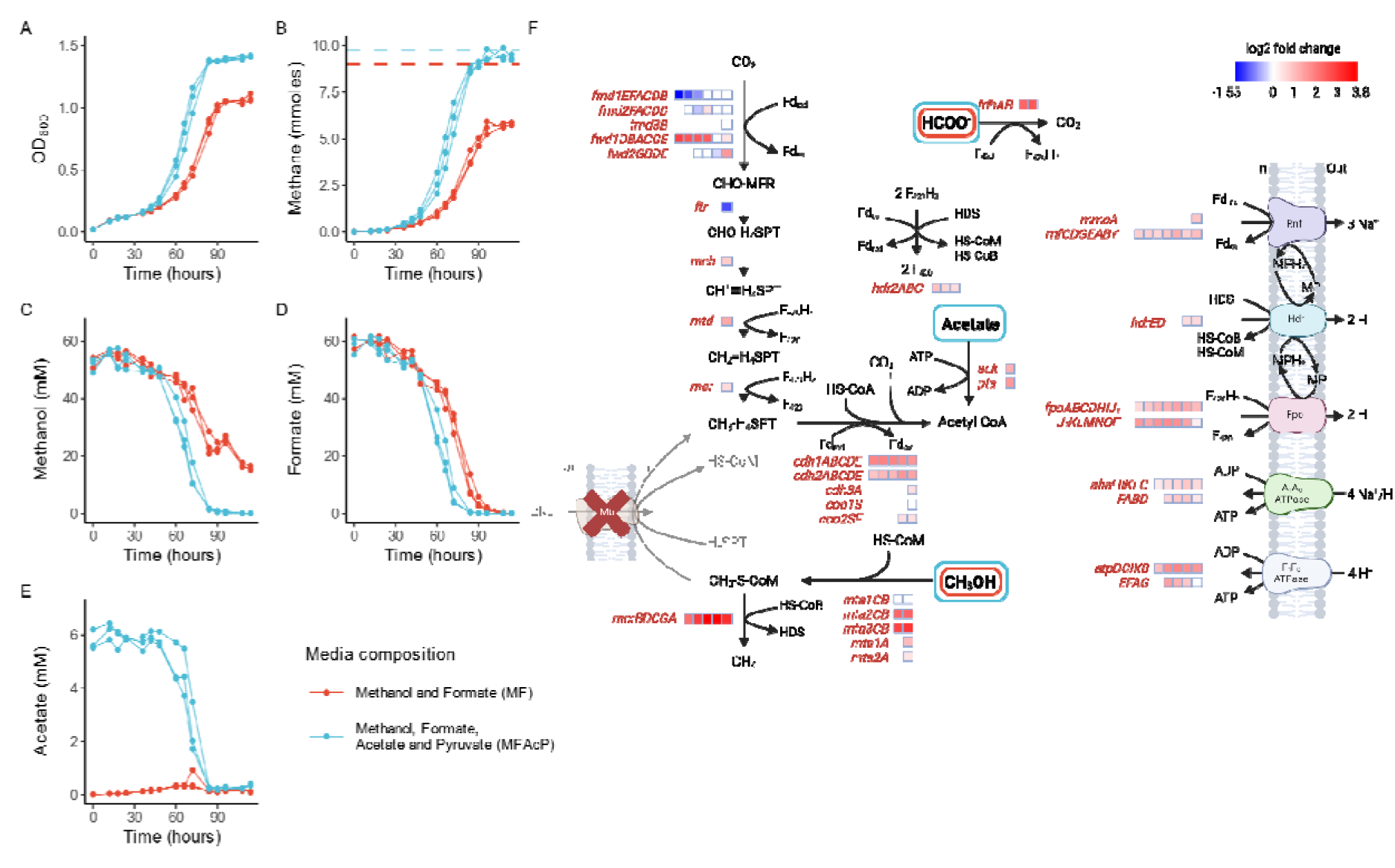
Growth of *M. acetivorans* JBA01 with formate-dependent methyl-reducing methanogenesis. The strain was grown in three biological replicates in media containing methanol and formate only (MF, Red) and in media where acetate and pyruvate is also provided (MFAcP, Blue). All replicates are shown (A) OD_600_ (B) Methane level [mmoles] Dashed lines represent theoretical yield of methane (C) Methanol level [mM] (D) Formate level [mM] (E) Acetate level [mM] and (F) Methanogenesis pathway in *M. acetivorans* JBA01 with heatmap showing differential gene expression analysis comparing growth in MFAcP to growth in MF. Log2 fold change indicates the change in the expression level in MF media compared to MFAcP media. Abbreviations: ferredoxin (Fd), methanophenazine (MP), methanofuran (MFR), tetrahydrosarcinapterin (H_4_SPT). Media composition: High salt media with methanol, formate, acetate, and pyruvate (MFAcP), High salt media with methanol and formate (MF). Gene for enzymes: formylmethanofuran dehydrogenase (*fmd/fwd*), formyl transferase (*ftr*), methenyl-H_4_SPT cyclohydrolase (*mch*), methylene-H_4_SPT reductase (*mer*), methyl-H_4_SPT:HS-CoM methyltransferase (*mtr*), methyl-CoM reductase (*mcr*), heterodisulfide reductase (*hdr*), methanol:HS-CoM methyltransferase (*mta*), carbon dioxide dehydrogenase (*cdh*), acetate kinase (*ack*), phosphate acetyltransferase (*pta*), F_420_H_2_ dehydrogenase (*fpo*), *Rhodobacter* nitrogen fixation complex (*rnf*). For genes with multiple homologs, the Gene IDs can be found in SI Appendix text.

The strain was also able to grow in absence of acetate and pyruvate, in MF media containing only 60 mM methanol and formate each. The doubling time was ≈ 15 hours in MF media, which was longer when anabolic substrates were absent (Fig. 3A). Metabolite analysis showed that all formate was consumed by the end of growth, but 15.7 ± 0.8 mM (≈ 25%) methanol remained (Fig. 3C-D). Since *M. acetivorans* JBA01 grows in media that does not contain acetate and pyruvate, it must have redundancy for Fd_red_ generation that does not rely on methanol oxidation. The difference in observed stoichiometry of five formate molecules oxidized to reduce three methanol in contrast of expected one formate for one methanol for catabolism is most likely because of formate requirement in anabolic reactions (Fig. 3C-D).

The transcriptome was analyzed to understand how *M. acetivorans* JBA01 was able to regenerate Fd_red_ and in turn cellular carbon during formate-dependent methyl-reducing methanogenesis. Differential gene expression analysis was done comparing with or without exogenous acetate and pyruvate. 1814/4544 DEGs (p_adj_ < 0.01) showed significant upregulation or downregulation (|Log2FC| > 0.5) (SI Appendix Table S2 and Dataset S2) in absence of acetate and pyruvate. Firstly, the *fdh*, inserted in the *mtr* operon was upregulated in media without acetate and pyruvate. Along with *fdh, rnf* was significantly upregulated. All subunits of the *fpo* complex were significantly upregulated except for *fpoF* (Fig. 3F). As was the case before, 907 DEGs were annotated as predicted/conserved hypothetical proteins and 11 being transcriptional regulators (SI Appendix Dataset S2).

### Formate-dependent CO_2_-reducing methanogenesis

Formate-dependent CO_2_-reducing methanogenesis pathway has never been shown in Methanosarcinales (1). We integrated *fdhAB* in F_420_-reducing hydrogenase (*frh)* operon without disrupting *mtr* as it would be essential for this methanogenesis pathway. The strain obtained, *M. acetivorans* JBAF01, was subjected to adaptive laboratory evolution (ALE) to increase the fitness and to be able to grow faster solely on formate in HSF media (HSF = 120 mM formate only). The serial transfers of *M. acetivorans* JBAF01 were started from methanol media into rich formate media containing casamino acids along with 5 mM acetate and pyruvate and later transferred to HSF media. At the end of repeated serial transfers, strain *M. acetivorans* JBAF02 (new designation at the end of ALE) had shortened doubling time of methanogenesis from ≈ 80 hours to ≈ 11 hours when growing solely on formate (Fig. 4A, B, F). However, at the end of ALE, *M. acetivorans* JBAF02 had increased doubling time of methanogenesis and slower growth (doubling time ≈ 12 hours) on methanol compared to that of *M. acetivorans* WWM73 (doubling time ≈ 7 hours) (Fig. 4A, C; Fig. S3A, C). The biomass yield of JBAF02 growing on formate was 2.48 g total protein mol^-1^ CH_4_, which was highest than the other growth conditions investigated in this study (SI Appendix Table S3). The genomes of selected strains throughout ALE were sequenced and analyzed. One structural variation, 332-bp deletion in *fwdD1*, was observed in the genome. *fwdD1* encodes the subunit D of formylmethanofuran dehydrogenase responsible for reversible reduction of CO_2_ to formyl group which is then transferred to methanofuran. *M. acetivorans* encodes four homologs of formylmethanofuran dehydrogenase, two annotated as tungsten-dependent and the other two as molybdenum-dependent (25). To check whether deletion in *fwdD1* was the reason for faster growth, we recreated the deletion in *M. acetivorans* JBAF01 (strain before ALE), but this deletion failed to decrease the methane doubling time in *M. acetivorans* JBAF01 Δ*fwdD1* (SI Appendix Fig. S4). 5 SNPs were also present in *M. acetivorans* JBAF02 that were not present in *M. acetivorans* JBAF01, summarized in SI Appendix Table S4.

**Figure 4.**
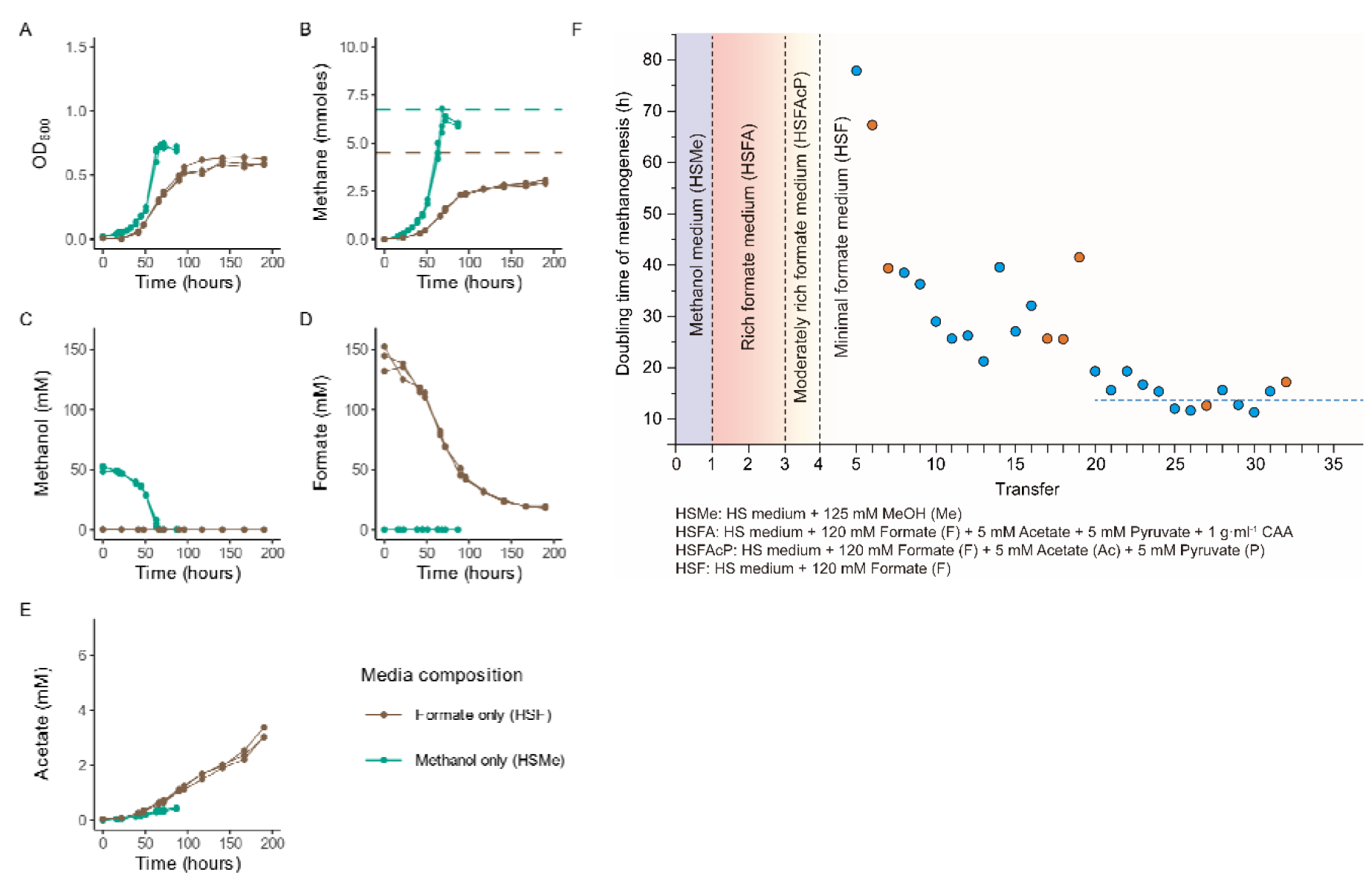
Growth of *M. acetivorans* JBAF02 with formate-dependent CO_2_-reducing methanogenesis and methylotrophic. The strain was grown in media containing formate as the sole carbon source (HSF, Brown) and in media containing methanol as the sole carbon source (HSMe, Green). (A) OD_600_ (B) Methane level [mmoles] Dashed lines represent theoretical yield of methane for each condition(C) Methanol level [mM] (D) Formate level [mM] (E) Acetate level [mM] and (F) Doubling time of methanogenesis during ALE. The ALE was done as described in the methods. Orange dots represent the transfer strains selected for genome sequencing.

When growing solely on formate, *M. acetivorans* JBAF02 did not utilize all available formate in the media. Approx 11 mM formate (8% of added formate) remained at the end of growth (Fig. 4D), unlike formate-dependent methyl-reducing methanogenesis of *M. acetivorans* JBA01 where all available formate was utilized (Fig. 3D). *M. acetivorans* JBAF02 also produced over 3 mM acetate when growing solely on formate (Fig. 4E).

Although JBAF02 exhibits similar metabolism as cytochrome-free methanogens, cytochromes are still integral to the growth of *M. acetivorans*. This was confirmed by addition of diphenyleneiodonium chloride (DPI), a methanophenazine analogue that has been shown to block electron transport in the membrane of *M. mazei*, to the media (SI Appendix Table S5) (26). DPI inhibited the growth of *M. acetivorans* JBAF02 but did not do the same for *M. maripaludis*, showing that electron transport through membrane was still crucial for growth.

Transcriptomic analysis was conducted to understand how formate is incorporated and how Fd_red_ is generated. Differential gene expression analysis was done to compare growth on formate with that on methanol of *M. acetivorans* JBAF02 (SI Appendix Fig S5). A total of 1055 DEGs (p_adj_ < 0.01) showed significant upregulation or downregulation (|Log2FC| > 0.5) (SI Appendix Table S2 and Dataset S3).

As was the case with *M. acetivorans* WWM73, expression of *fdhD* stayed relatively unchanged, only slightly downregulated (Log2FC = -0.21) in presence of formate. *ma4008* (*aceP*) was upregulated, again possibly for acetate efflux. In this analysis, *ma4393* and *ma0103*, the other two uncharacterized AceTr family proteins, were also differentially expressed. *ma4393* was downregulated (Log2FC = -2.43) in presence of formate whereas *ma0103* was slightly upregulated (Log2FC = 0.43). No significant change was seen in *rnf* or *fpo* transcription. *hdrA1* (Log2FC = 0.55) and *hdrB2C2* (Log2FC ≈ 0.55) were significantly upregulated in presence of formate compared to methanol. As was the case before, many DEGs (497) were annotated as predicted/conserved hypothetical proteins and 15 being transcriptional regulators (SI Appendix Dataset S3).

Differential gene expression analysis was also done between *M. acetivorans* WWM73 growing on methanol with *M. acetivorans* JBAF02 on methanol as well as formate (SI Appendix Fig S6 and S7 and Dataset S4 and S5) even though the genetic makeup was different. This comparison was done to get a hint about differential expression of any energy conservation pathway in *M. acetivorans* JBAF02 when growing on formate or methanol compared to unedited strain. Notable differential expression was seen F_1_F_0_ ATPase and methanol-specific methyltransferases.

## Discussion

### Newly engineered strains of *M. acetivorans*

*M. acetivorans* JBAF02 grows by a metabolism novel to the order and to cytochrome-containing methanogens. Formate-dependent CO_2_-reducing methanogenesis in *M. acetivorans* JBAF02 highlights the strains capability to produce Fd_red_ required for CO_2_ reduction and anabolism from F_420_H_2_ without using H_2_ as an intermediate. Although FdhAB enabled CO_2_ reduction to methane, the process was slow in the beginning. ALE increased the fitness of the strain on formate but *M. acetivorans* JBAF02 had slower growth on methanol comparable to that of *M. acetivorans* WWM73, which could be due to the significant downregulation of the genes of the major methyltransferases, *mtaC1* and *mtaC2B2* (Fig. S6).

*M. acetivorans* JBA01 became capable of formate-dependent methyl-reducing methanogenesis with heterologous expression of formate dehydrogenase and disruption of *mtr*. Outside *Methanosarcina*, a similar pathway is naturally occurring only in the members of *Methanonatronarchaea. Methanonatronarchaeum thermophilum* uses a membrane bound formate dehydrogenase that directly reduces methanophenazine using formate. *M. thermophilum* also requires exogenous acetate to be provided for growth and even then, its doubling time on methanol and formate was slower (≈ 49 hours) than that of *M. acetivorans* JBA01 (≈ 15 hours) (14).

The difference in formate utilization threshold of *M. acetivorans* JBA01 and JBAF02 is likely due to the difference in the energy metabolism of the two strains. The calculated range of max ATP yield, given the formate threshold, is 1 to 1.6 ATP mol^-1^ CH_4_ for *M. acetivorans* JBAF02 when growing on formate (not accounting for possible substrate level phosphorylation in acetate production, calculations in SI Appendix text).

An emerging acetogenic potential is also seen in presence of formate where *M. acetivorans* WWM73 produced approx. 1.5 mM acetate and *M. acetivorans* JBAF02 produced approx. 3.6 mM acetate. Acetogenic potential of *M. acetivorans* has been discussed when growing on CO where over 75% of supplied CO is converted to acetate (27).

In all transcriptomic comparisons, the expression levels of *mrp* Na^+^/H^+^ antiporter remained constant except when *M. acetivorans* JBA01 is cultured on MF compared to MFAcP. The Mrp complex plays a critical role in efficient ATP synthesis and promoting optimal growth in *M. acetivorans* when utilizing acetate, especially when the acetate level is low (28). The observed reduction in *mrp* transcription levels in *M. acetivorans* JBA01 during growth on MFAcP could be attributed to a global downregulation of energy-related genes, including *mcr*, ATP synthases, and cytochromes. This adjustment is probably a response to an abundance of nutrients in the environment, reducing the need for enhanced energy conservation mechanisms.

*M. acetivorans* harbors two ATP synthases: the bacterial F_1_F_0_ ATPase, which utilizes a proton gradient for ATP synthesis, and the archaeal A_1_A_0_ ATPase, capable of using both proton and sodium gradients. Notably, the bacterial F_1_F_0_ ATPase is not essential for *M. acetivorans* growth (29). The expression level of F_1_F_0_ ATPase is significantly upregulated during formate-involved metabolism in the absence of the anabolic substrates. This observation aligns with the generated energy conservation mechanisms in formate-dependent methylotrophy and formate-dependent CO_2_ reduction, which contrast with traditional methylotrophy by producing a negative/no sodium gradient and a positive proton gradient (SI Appendix Table S6). Consequently, F_1_F_0_ ATPases likely contribute to proton translocation for ATP synthesis and pH homeostasis within these specific metabolic pathways.

The plasticity in energy metabolism of *M. acetivorans* is highlighted by the different options to generate Fd_red_ during formate-dependent growth. The high hydrogen threshold required for cytochrome-containing methanogens has been hypothesized to prevent them from utilizing formate via hydrogen cycling (1, 30). As the energy metabolism of *M. acetivorans* does not rely on hydrogen cycling, it is able to utilize formate (21). In the methanogenesis pathways of *M. acetivorans* JBA01 (Fig. 1D) and *M. acetivorans* JBAF02 (Fig. 1E), the cell can generate Fd_red_ only from F_420_H_2_. The possible pathways are summarized in SI Appendix Figure S8. (1) Membrane-bound Rnf running in reverse (31, 32), (2) the HdrA2B2C2 enzyme (33), (3) the HdrA1B1C1 forming complex with FdhAB (34, 35), and (4) the membrane bound Fpo without FpoF (36). All four pathways may also sum up to the observed effect. Detailed support for each hypothesis is discussed in SI Appendix text.

### Native capability of *M. acetivorans* towards formate utilization

The genome of *M. acetivorans* encodes a *fdhD* gene which is a formate dehydrogenase family accessory protein catalyzing the sulfur transfer from *L*-cysteine desulfurase (*IscS)* to formate dehydrogenase. In *Escherichia coli, fdhD* has been shown to be essential for formate dehydrogenase activity (37). The presence and expression of this gene could be as important in *M. acetivorans* to make it amenable for growth and methanogenesis from formate. As formate does not change the expression of this gene significantly, the corresponding accessory protein might be essential only in small amounts.

There are no formate transporters annotated in the genome of *M. acetivorans* (38). Recent studies have shown the importance of formate transporters for growth on formate. In *Methanococci*, growth on formate was dependent not only on the formate dehydrogenase genes but also on the presence of a formate transporter FdhC (39). As there is no *fdhC* homolog found in the genome of *M. acetivorans*, the microbe is expected to use another transporter for formate intake. AceTr family proteins are shown to support formate transport in *Saccharomyces cerevisiae* (40). Three genes belonging to the AceTr family are present in the genome of *M. acetivorans* out of which only one, *ma4008*, has been characterized and shown to be acetate specific (23). It is upregulated in the presence of formate for acetate efflux from the (41). *ma4393* and *ma0103* are uncharacterized and could transport formate. In *M. acetivorans* JBAF02, *ma4393* was significantly downregulated in presence of formate whereas *ma0103* was slightly upregulated. Characterizing *ma4393* and *ma0103* could enable us to identify the unannotated formate transporter in *M. acetivorans*.

### Distribution of FdhA in Methanosarcinales

The ubiquitous presence of FdhA and its multiple acquisitions in *Methanotrichaceae* suggests that this enzyme has a key role in these methanogens. *Methanothrix soehngenii* cleaves formate into H_2_ and CO_2_ (42), demonstrating that *fdhA* is expressed, but none of the described *Methanothrix* species can use formate for methanogenesis (43, 44). Similarly, *M. barkeri* has been shown to have formate dehydrogenase activity yet attempts to grow *M. barkeri* solely on formate have failed in the past (18, 19). The role of the formate dehydrogenase in these ecologically important methanogens needs to be clarified.

FdhA may have been present in the last common ancestor of the Methanosarcinales and lost after the divergence between the Methanotrichaceae and other Methanosarcinales. Then FdhA was re-acquired in some Methanosarcinaceae, including *M. barkeri*, through HGT. Multiple acquisition of this gene in Methanosarcinaceae highlights the metabolic flexibility of members of the family that have accommodated several times the utilization of formate in their metabolic pathways. This was illustrated here by the heterologous expression of FdhAB from *M. barkeri* in *M. acetivorans* and showing formate-dependent methanogenesis in *M. barkeri* as well.

*M. acetivorans* might have lost the formate dehydrogenase but has retained the accessory machinery for enabling formate utilization and the redundancy in energy metabolism to readily incorporate formate. In the presence of formate over 800 genes were differentially expressed in *M. acetivorans* WWM73. These DEGs included genes for methanogenesis and energy metabolism. *M. acetivorans* therefore senses and adapts to the presence of formate. These transcriptomic changes may be a native response to the formate produced when growing on CO by *M. acetivorans* (27).

Formate has been shown to be abiotically formed near hydrothermal vents and was present in early earth (39). Methanogenesis has been hypothesized to have originated methylotrophically by using H_2_ as the reductant prior to evolution of *mtr* that enabled disproportionation of methyl groups (45). Even if not in the origin of methanogenesis but somewhere along the evolutionary line, a methanogen would have existed, ancestor to *M. acetivorans*, capable of reducing methylated substrates with formate. The question remains as to why this ability was lost. Was it competition for formate or lack of fitness due to formate? Also, why is this metabolism so rare in nature?

Our efforts to include *fdhAB* along with disruption of *mtr* and ALE were the first steps on the path to enable formate-dependent methanogenesis in *M. acetivorans*. However, it was only made possible by *M. acetivorans* meeting us halfway with its flexibility and redundancy in the energy metabolism. Unlocking formate as an electron donor in the metabolism of the *M. acetivorans*, we have made an already versatile methanogen even more versatile. This allows for a previously unavailable metabolic engineering pathway to be used for new ways of sustainable C1-valorization. Enabling formate dependent methanogenesis in *M. acetivorans* is a good example of Nature (the plasticity of metabolism towards formate) and Nurture (our genetic engineering efforts with ALE) working in tandem towards a common goal.

## Materials and Methods

### Microbiological and molecular methods

Lysogenic broth containing 50 mg L^-1^ ampicillin was used for plasmid construction in *E. coli* NEB5α. *M. acetivorans* was cultured in a high-salt medium tailored to the specific requirements of the experiment (46). Depending on the experimental conditions, substrates were added as follows: 60 mM methanol alone; 60 mM methanol combined with 60 mM formate; a mixture of 60 mM methanol, 60 mM formate, along with 5 mM each of acetate and pyruvate; or 120 mM formate alone. Cultures were grown at 37°C with the gas phase consisting of 50% N_2_/ 20% CO_2_/ 30% of 1% H_2_S at 1 bar. The optical density of the cultures was tracked using Eppendorf BioPhotometer plus spectrophotometer. Diphenyleneiodonium chloride (Sigma-Aldrich) was dissolved in dimethyl sulfoxide and added where mentioned to the media at a final concentration of 20 µM. Plasmid construction was carried out according to standard protocols. Detailed information is available in SI Appendix text. Liposome-mediated methods and polyethylene glycol-mediated methods were used to transform *M. acetivorans* (12). An optimized CRISPR/Cas9 genome editing toolbox was used to delete genes (11, 13). The media composition, strains, primers and gRNA, and plasmids used in this study were listed in SI Appendix Table S7, S8, S9, and S10 respectively. Detailed methods for strain construction are included in SI Appendix text.

### Quantification of formate dehydrogenase activity

Formate dehydrogenase activity was determined anaerobically by tracking formate-dependent benzylviologen reduction at 578 nm using cleared cell-free lysates. Detailed methods in SI Appendix text.

### Metabolite analyses

Methanol, formate, and acetate were quantified using ^1^H-NMR spectroscopy. Growth medium samples were complemented with 10% D_2_O and 0.10% 1,4-dioxane as an internal standard for quantification. The methane level in the headspace gas sample was measured using a GC-FID (Agilent HP 6890 Gas Chromatograph, Hewlett-Packard), equipped with an HP-AL/KCL column (length, 50 m; diameter, 0.320 mm, thickness, 8 µm). The headspace gas samples were injected with a Gastight 1700 SampleLock Syringe (100 µL, PN81056) (Hamilton). Detailed instrument methods for ^1^H-NMR and GC-FID in SI Appendix text.

### Adaptive laboratory evolution

*M. acetivorans* JBAF01 was initially cultivated in HSMe medium supplemented with 125 mM methanol (Transfer 1, T1). Subsequently, cells were washed twice using HS medium and then transferred (T2) to 10 mL of HSFA medium, which contained 120 mM formate, 5 mM each of acetate and pyruvate, and 1 g L^-1^ casamino acids. After four weeks, 1 mL from T2 culture was transferred (T3) to another 10 mL of HSFA medium. After 23 days, 1 mL from the T3 culture was transferred (T4) to 10 mL of HSFAcP medium, containing 120 mM formate, 5 mM each of acetate and pyruvate. After 16 days, 1 mL from the T4 culture was transferred (T5) to 10 mL of HSF medium. Gas chromatography (GC) measurements were conducted from the T5 culture onward, ranging from every four days to daily, depending on the rate of methanogenesis. Once the exponential phase of methanogenesis finished, 1 mL of the culture was transferred to a fresh 10 mL of HSF medium. From T24 forward, the volume of inoculum was reduced to 200 µL to accommodate the increased growth rate.

### Nucleic acid isolation, sequencing, and analysis

The genomic DNA was extracted with Wizard Genomic DNA Purification Kit (Promega). The Microbial WGS was performed using Illumina NovaSeq 6000 by Novogene (UK) Company. The resultant data was filtered using Trimmomatic (47), aligned using BWA-mem2 (48) to *M. acetivorans* C2A genome (38) (Accession no AE010299) and analyzed for SNPs and SVs using Samtools (49) and BreakDancer (50). For transcriptomics, the cultures were grown in triplicates to an OD of 0.3-0.4 and pelleted in RNAprotect. RNA was isolated using Qiagen RNA isolation kit. The total RNA was sent to Novogene for library prep and sequencing. The resultant data was filtered using Trimmomatic and analyzed using Geneious Prime 2023.1.2 Workbench. For genome reference for transcriptomics, *M. acetivorans* C2A genome (Accession no AE010299) was modified to have 4 extra sequences reflecting CDS of fdhAB, tetR and phi31. The differential gene expression analysis was done using DESEQ2 (51) in Geneious Prime 2023.1.2.

### Phylogenetic analysis

We searched the homologues of FdhA using HMMsearch from HMMER v3.3.2 suite (52), using an HMM profile built from an alignment of several FdhA sequences. The sequences presenting a e-value < 0.01 have been selected and aligned using MAFFT v7.490 (53) (default parameters). The alignment has been trimmed using BMGE v1.12 (54) (-b 1 -w 1 -h 0.95 -m BLOSUM30 options) and the trimmed alignment has been used to infer a phylogenetic tree using FASTTREE v2.1.10 (55) (LG+G4). We then delineated the FdhA subfamily among the big homologue family using the topology, branch length and sequence length.

For the phylogenetic analysis, the sequences have been sampled using a selection of 401 bacteria (8) and of 525 archaea (457 archaea corresponding to 1-15 genomes per major clade and 68 genomes of Methanosarcinales). We also added sequence of *M. mazei* identified by a BLASTp on NCBI, 2 sequences of other Methanosarcinales and 2 sequences of Methanoflorentaceae that were not present in the initial database. The sequences have been realigned using MAFFT-linsi (53), and the resulting alignment has been manually curated. The alignment has then been trimmed using BMGE (54) (-b 1 -w 1 -h 0.95 -m BLOSUM30 options) and the trimmed alignment has been used to infer a phylogenetic tree using IQ-TREE v1.6.12 (56), using the best suited model according to the BIC (LG+R10). The robustness of the branches has been assessed by 1,000 ultrafast bootstrap replicates. The figure has been generated using iTOL (57).

## Data availability statement

All raw reads for whole-genome sequencing and transcriptomics have been uploaded to SRA under the Bioproject PRJNA1085037. All other study data is available in the article and/or SI Appendix.

## Acknowledgments

We thank Prof. Alfred Spormann, Dr. Boyang Ji and Dr. Christian Molina for helpful discussions. This research has received financial support from the NovoNordisk foundation (Grant no NNF19OC0054329 to S.S. and grant no NNF20OC0065032 to J.B.) and the Research Council of Finland (Grant no 329510 to S.S.). The authors thank Prof. Metcalf (University of Illinois, USA) for providing the strain WWM73. The authors acknowledge the Aalto University Raw Materials Research Infrastructures and Bioeconomy facilities.

