## supplementary figures tables for "Nature AND Nurture: Enabling formate-dependent growth in *Methanosarcina acetivorans*"

**This PDF file includes:**

Supporting text

Figures S1 to S8

Tables S1 to S10

Legends for Datasets S1 to S5

SI References

**Other supporting materials for this manuscript include the following:**

Datasets S1 to S5

Supporting Text

Estimation of the free energy available after growth of strain JBAF02 on formate.

Free energy for formatotrophic methanogenesis at standard conditions:

4 HCOO^-^ + 4 H^+^ = 3 CO_2_ + CH_4_ + 2 H_2_O	ΔG° = -304.0 kJ mol^-1^, ΔG°’ = -144.4 kJ mol^-1^

The free energy (ΔG) is calculated according to the following formula:

ΔG = ΔG° + RT ln(K)

For simplicity, we provide the calculation of the thermodynamics at T = 298 K (25 °C, standard conditions), even though the incubations were at 37 °C. This estimation gives a value (error) that is ca. 1.5 kJ mol^-1^ lower than if calculated at 310 K (37 °C, the actual incubation temperature).

K is defined as follows:

K = [CO_2_]^3^×[ CH_4_] / ([HCOO^-^]^4^×[H^+^]^4^)

Concentrations of products and residual substrates after incubation of JBAF02 with formate:

| **Species** | **Concentration** | **Comment** |
| --- | --- | --- |
| Formate (HCOO^-^) | 0.0115 M | Measured via ^1^H-NMR spectroscopy |
| Protons (H^+^) | 1.58 * 10^-8^ M | Measured pH = 7.8 |
| Methane (CH_4_) | 0.2 bar | Measured via GC |
| Carbon dioxide (CO_2_) | 0.2 – 1.2 bar | Estimation |

The CO_2_ partial pressure could not be directly measured. If all CO_2_ derived from the consumed formate would end up in the headspace, a partial pressure of 1.2 bar would be obtained, which serves as the upper limit. When comparing methanogenesis by JBAF02 on formate vs. on methanol, we observed that growth on methanol leads to a much stronger pressure build-up compared to formatotropic methanogenesis, even though in both cases the same amount of gas would be expected. The CO_2_ formed from formate is putatively absorbed partly in the medium due to the pH increase. We therefore estimate 0.2 bar CO_2_ as the lower limit, which corresponds to the initial partial pressure if all newly formed CO_2_ would stay in solution.

Results of the thermodynamic calculations:

For a CO_2_ partial pressure of 0.2 bar:

K = 1.45 * 10^36, ΔG = -97.7 kJ mol^-1^

For a CO_2_ partial pressure of 1.2 bar:

K = 3.13 * 10^38, ΔG = -84.4 kJ mol^-1^

Taking uncertainties of other values into account (e.g. pH measurement, influence of temperature), we conclude that the free energy is between -100 kJ mol^-1^ and -80 kJ mol^-1^.

Assuming the calculated range of free energy and the ΔG for ATP formation is 60 to 80 kJ mol^-1^(1), we get can ATP yield of 1 – 1.67 ATP mol^-1^ CH_4._

Different options to generate Fd_red_ during formate-dependent growth in *M. acetivorans*

SI Appendix Fig S8A - Membrane-bound Rnf running in reverse

The *Rhodobacter* nitrogen fixation (Rnf) complex couples the oxidation of Fd_red_ to reduction of methanophenazine and pumps out 3 Na^+^ during this process in *M. acetivorans* (2, 3). The reduced methanophenazine then reduces the heterodisulfide (HDS) of coenzyme M and coenzyme B via the membrane-bound enzyme HdrED that pumps out additional ions in the form of H^+^. In other microbes, Rnf has shown to have the activity of Fd:NADH oxidoreductase (4). The complex was also shown to work in reverse as well in in vitro experiments. Using Na^+^ gradient, it was able to catalyse endergonic reduction of Fd_ox_ using NADH (4). As Rnf is reversible, it is the most likely solution for *M. acetivorans* to produce Fd_red_ when growing on formate.

SI Appendix Fig S8B - Electron bifurcation via HdrA2B2C2

HdrA2B2C2 has been shown to have cytoplasmic Fd:HDS oxidoreductase activity with F_420_H_2_ as the electron donor(5). The complex has been shown to use electron bifurcation to reduce HDS and Fd_ox_ by electrons obtained from F_420_H_2_. Given the abundance of F_420_H_2_ in the strains designed in this study, it is likely that this protein could help with generation of Fd_red_

$$2\text{ F}_{420}\text{H}_{2}+\text{Fd}\text{ox}+\text{CoM}\text{-S}\text{-}\text{S-}\text{CoB}\to\mathrm{Fd}_{\mathrm{red}}\text{+ HS-CoM}+\text{HS-CoB}+2 \text{F}_{\text{420}}$$

SI Appendix Fig S8C - Electron bifurcation via HdrA1B1C1 and FdhAB complex

HdrABC and FdhAB have been shown to form a multienzyme complex for electron bifurcation to reduce HDS and Fd_ox_ concurrently using formate in *Methanococcus maripaludis*. The multienzyme complex consists of HdrABC, FdhAB and MvhD(6). There is no *mvhD* annotated gene in *M. acetivorans* but the HdrA of *M. acetivorans* has been shown to have a fused MvhD domain (5). Similar to *Methanomicrobiales*, the HdrABC from *M. acetivorans* could form a complex with FdhAB to enable electron bifurcation from formate to HDS and Fd_ox_.

SI Appendix Fig S8D - “Headless Fpo”- Fpo complex without FpoF

*Methanosaeta thermophilum* genome encodes the protein complex Fpo without the FpoF subunit. This “headless” Fpo, lacking the F_420_H_2_ oxidizing subunit FpoF, has been hypothesized to accept electrons from Fd_red_. The support for this hypothesis comes from homologues in cyanobacteria and chloroplasts that do not contain FpoF homologue use Fd_red_ as electron donor (2). Similar to the hypothesis that Rnf could be reversible, it is possible that “headless” Fpo complex runs in reverse to catalyse generation of Fd_red_ using proton gradient and reduced methanophenazine.

Strain construction

The construction of strain JBA01 was achieved by introducing the recombinant fragment "mtr::fdh P1P10" into *Methanosarcina acetivorans* WWM73, followed by screening on MFAcP plates supplemented with 1 g L^-1^ casamino acids and 2 µg mL^-1^ puromycin. The "mtr::fdh P1P10" fragment was generated through overlap PCR by concatenating fragments "mtr::fdh P1P2", "mtr::fdh P3P4", "mtr::fdh P5P6", "mtr::fdh P7P8", and "mtr::fdh P9P10". Fragments "mtr::fdh P1P2" and "mtr::fdh P9P10" were amplified using the *M. acetivorans* WWM73 genome as a template, whereas "mtr::fdh P3P4" and "mtr::fdh P5P6" utilized the *Methanosarcina barkeri* WWM155 genome. The "mtr::fdh P7P8" fragment was amplified from pM001.

The excision of the *pac-hpt* selection marker from the strain was achieved by introducing the "mtr::fdh P11P14" fragment into the previously mentioned strain. This fragment includes approximately 1.1 kb of homologous recombination regions upstream and downstream of the *pac-hpt* cassette. Following this introduction, counterselection was performed on MFAcP plates containing 20 µg mL^-1^ 8-aza-2,6-diaminopurine (8ADP).

Strain *M. acetivorans* JBAF01 was engineered by transforming PvuI-linearized pfdh V3 into *M. acetivorans* WWM73, followed by screening on HSMe plates with 2 µg mL^-1^ puromycin. The pfdh V3 plasmid was assembled using Gibson assembly, incorporating fragments "frh::fdh P15P16", "frh::fdh P17P18", "frh::fdh P19P20", "frh::fdh P21P22", "frh::fdh P23P24", "frh::fdh P25P26", "frh::fdh P27P28", and the ApaI and SacI-digested p425GPD vector.

The *pac-hpt* marker was excised through counterselection on HSMe plates containing 20 µg mL^-1^ 8ADP.

The rest of the strains were constructed by transfroming the corresponding plasmids.

The assembly of additional plasmids was performed using Gibson cloning and Golden Gate cloning techniques, adhering to standard plasmid construction protocols (7, 8).

Gene IDs of genes with multiple homologs

*fmd1EFACDB	ma0304-0309*

*fmd2FACDB	ma4174-4179*

*fmd3B	ma1241*

*fwd1DBACGE	ma0835-0832, 0671, 0381*

*fwd2GBD	ma2877-2879*

*hdr2ABC	ma2868, 4237-4236*

*hdrED	ma0687-0688*

*cdh1ABCDE	ma1016-1011*

*cdh2ABCDE	ma3860-3862,3864-3865*

*cdh3A	ma4399*

*coo1S	ma1309*

*coo2SF	ma3282-3283*

**Detailed materials and methods**

**Quantification of formate dehydrogenase activity.** 10 ml cultures of *M. acetivorans*, propagated on methanol, were harvested by centrifugation (5,400 g) for 10 min. Cleared cell-free lysates (CCL) were prepared by resuspending the pellets in 1 ml potassium phosphate buffer (KP, 50 mM, pH 7.2) and incubating for 30 min on ice before centrifuging (5,400 g, 10 min) again.

Fdh activity was determined anaerobically in stoppered cuvettes (flushed with N2) from the supernatant by following formate-dependent benzylviologen (BV) reduction (at 578 nm for 5 min); 1 ml assays consisted of 750 µl KP, 100 µl 1M formate, 100 µl (BV 10 mM in KP); before starting the assay with 50 µl CCL, BV was slightly pre-reduced with 10 µl 100 mM Na-dithionite (until light blue); stock solutions were made anaerobic by repeated gas/vacuum cycles in stoppered vials; formate-independent BV reduction of the extract was subtracted from the values to account for unspecific oxidoreductase activity.

Specific Fdh activity was calculated using an absorption coefficient for BV of 8.65 mM^-1^ cm^-1^ (9) and is given in U (1 µmol BV reduced per min) per mg protein. All protein quantification was done using the Pierce Kit and employing the method of Bradford (10).

**Metabolite analysis.** The ^1^H-NMR spectroscopy measurements were performed at 298 K on a 400 MHz Bruker Avance III spectrometer equipped with a 5 mm room temperature BBFO probe. Data acquisition was done using the Bruker TopSpin 3.6.2 software. The spectra were measured with a repetition time of 9.1s over 16 scans using a 1D NOESY method with presaturation during relaxation delay and mixing time. The chemical shifts were calibrated on the signal of dioxane, set to 3.70 ppm. The integration values of peaks at 8.39 ppm for formate (s), 3.30 ppm for methanol (s) and 1.57 ppm for acetate (s) used for quantification. The GC-FID was setup with the temperature of the front inlet set to 200 °C, that of the column oven to 40 °C (isothermal) and that of front detector to 250 °C. The pressure of the front inlet was set to 1.45 bar, and the total flow of helium was set to 15.2 mL min^-1^. The mobile phase was hydrogen and synthetic air at a flow rate of 35 mL min^-1^ and 350 mL min^-1^. The makeup flow was 26 mL min ^-1^ for helium.

**Database for phylogenetic analysis.** The database used to search for FdhA homologues contains 10,864 prokaryote proteomes from NCBI and has been assembled in a previous study (11). Several sampling steps were applied to assemble this database. First, we removed the redundancy based on taxonomic IDs. Second, we performed a whole genome comparison and used a clustering approach to group closest genomes, and then selected one representative genome per group. Third, we clustered the genomes by phylum/major clade and performed a clustering based on RpoB sequences. The representative genomes were selected using the NCBI completeness status, the NCBI representative status, the availability of annotation files and the number of proteins reviewed in Uniprot.

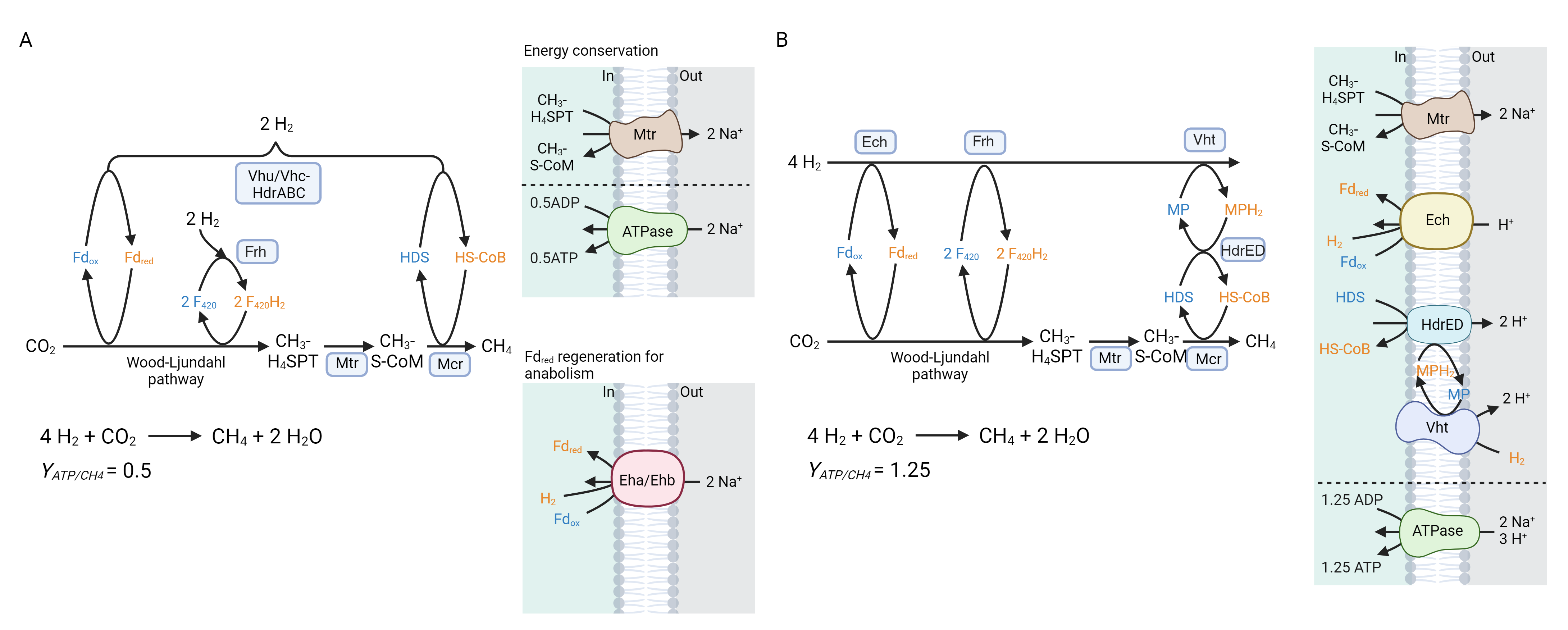

Fig. S1 Generation of Fd_red_ in H_2_-dependent CO_2_-reducing methanogenesis. (A) methanogens without cytochromes utilize electron bifurcation (HdrABC) as well as proton gradient generated via H_2_ (Eha/Ehb) to reduce different pools of ferredoxin required for CO_2_ reduction and anabolic reactions.(12) (B) methanogens with cytochromes utilize proton gradient generated via H_2_ to reduce ferredoxins required for both CO_2_ reduction and anabolic reactions (13). Similar strategy is used by microbes when growing on formate as well and as such methanogens with higher ATP yield (methanogens with cytochromes) require higher H_2_ threshold to grow on formate (as hypothesized in previous publication (14)). Chemical equations and ATP yield mentioned for each condition below the respective panel. Abbreviations: ferredoxin (Fd), methanophenazine (MP), F_420_-reducing hydrogenase (Frh), F_420_-non reducing hydrogenase (Vhu/Vhc), methanophenazine-reducing hydrogenase (Vht), energy converting hydrogenase (Eha/Ehb/Ech), heterodisulfide reductase (Hdr), methyl-coenzyme M reductase (Mcr), methyl-H_4_SPT:HS-CoM methyltransferase (Mtr).

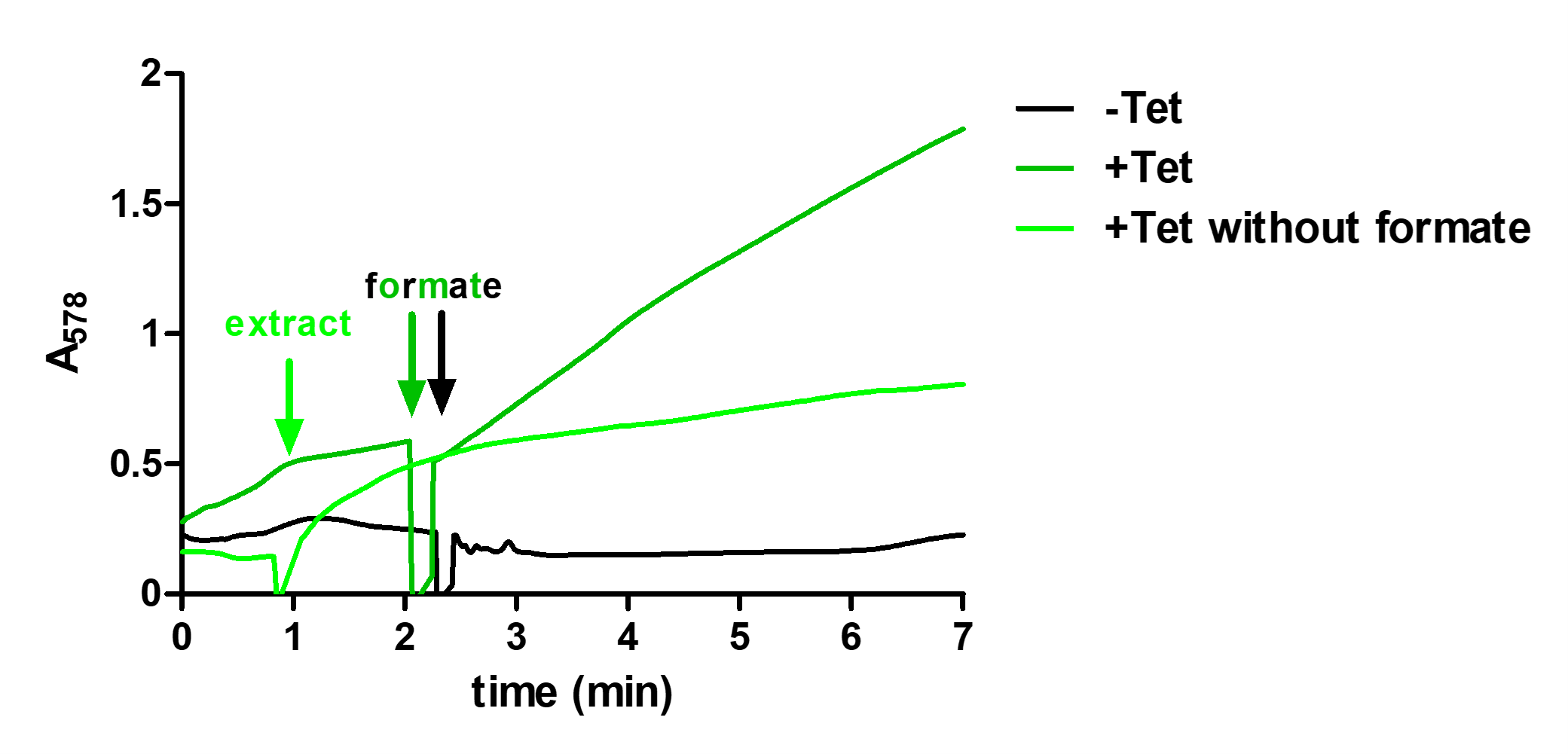

Fig. S2 *In vitro* characterization of FdhAB from *M. barkeri*. Reduction of benzyl viologen was measured. Dark green line indicates condition where formate was added along with expression of FdhAB. Light green line indicates condition where FdhAB was expressed without formate. Black line indicates where neither formate is added nor FdhAB was expressed. The activity measured was 0.72 U mg^-1^ total protein.

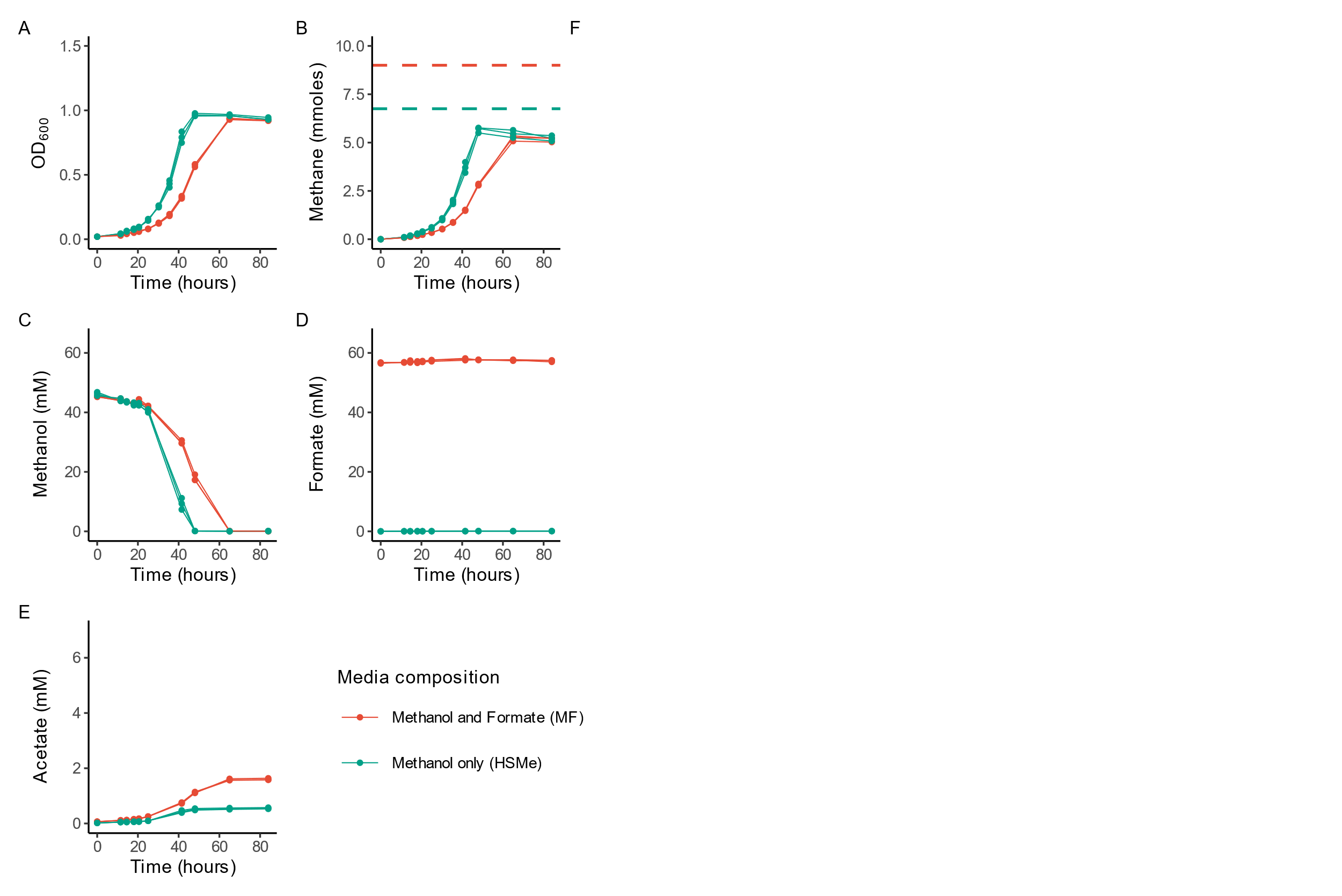

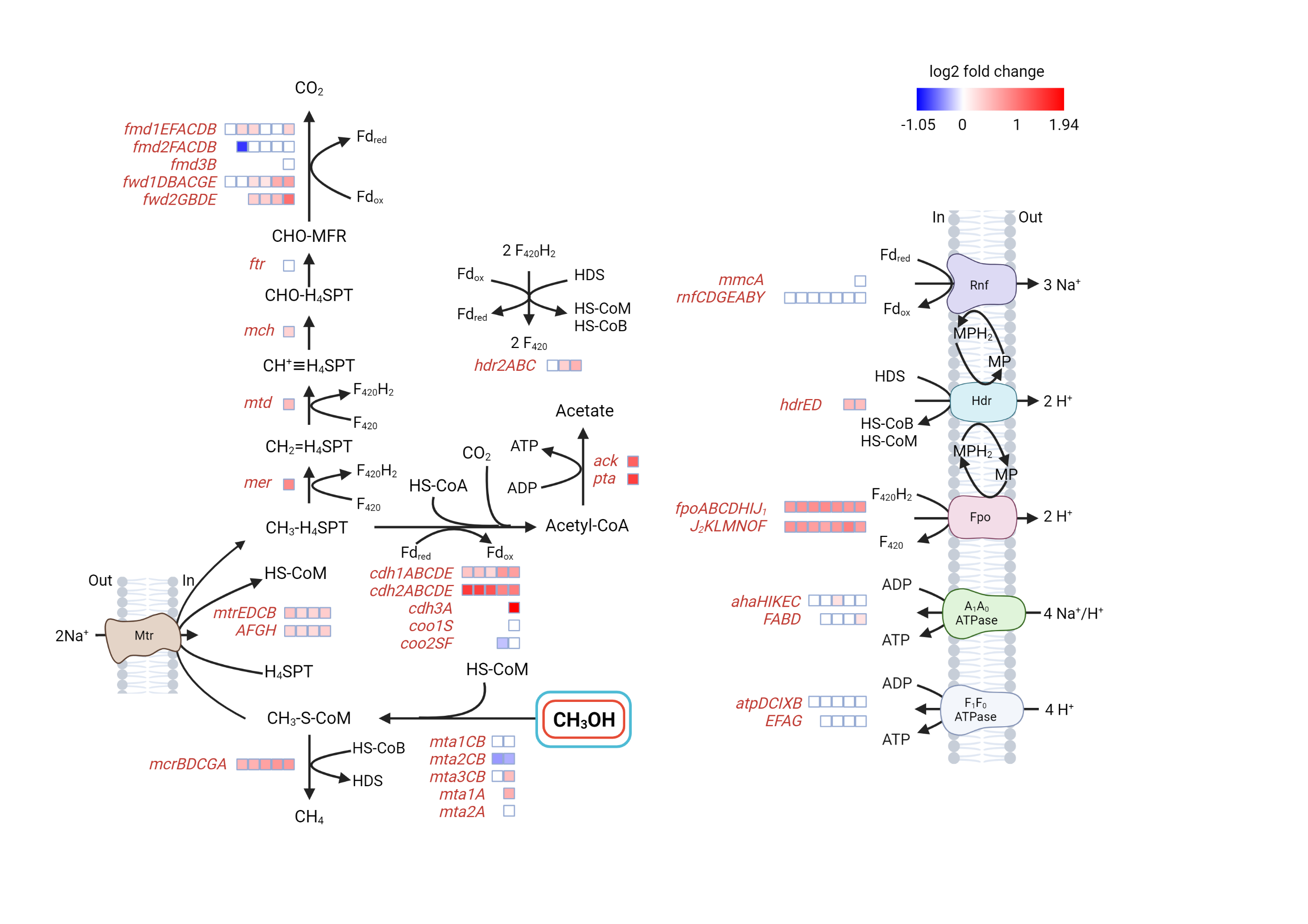

Fig. S3 Influence of formate on growth of *M. acetivorans* WWM73. The strain was grown in media containing methanol as the sole carbon source (HSMe, Green) and in media where formate is also provided (MF, Red). (A) OD_600_ (B) Methane level [mmoles] Dashed lines represent theoretical yield of methane for methanol disproportionation (Green) and for a hypothetical scenario where all methanol could be reduced to methane using electrons from formate(Red) (C) Methanol level [mM] (D) Formate level [mM] (E) Acetate level [mM] and (F) Methanogenesis pathway in *M. acetivorans* WWM73 with heatmap showing differential gene expression analysis comparing growth in HSMe to growth in MF. Log2 fold change indicates the change in the expression level in MF media compared to HSMe media. Abbreviations: ferredoxin (Fd), methanophenazine (MP), methanofuran (MFR), tetrahydrosarcinapterin (H_4_SPT). Media composition: High salt media with methanol (HSMe), High salt media with methanol and formate (MF). Genes: formylmethanofuran dehydrogenase (*fmd*/*fwd*), formyl transferase (*ftr*), methenyl-H_4_SPT cyclohydrolase (*mch*), methylene-H_4_SPT reductase (*mer*), methyl-H_4_SPT:HS-CoM methyltransferase (*mtr*), methyl-CoM reductase (*mcr*), heterodisulfide reductase (*hdr*), methanol:HS-CoM methyltransferase (*mta*), carbon dioxide dehydrogenase (*cdh*), acetate kinase (*ack*), F_420_H_2_ dehydrogenase *(fpo)*. Growth curve and metabolite curves obtained from three independent cultures (all shown). Transcriptomics done on three biological replicates. For genes with multiple homologs, the Gene IDs can be found in SI Appendix text.

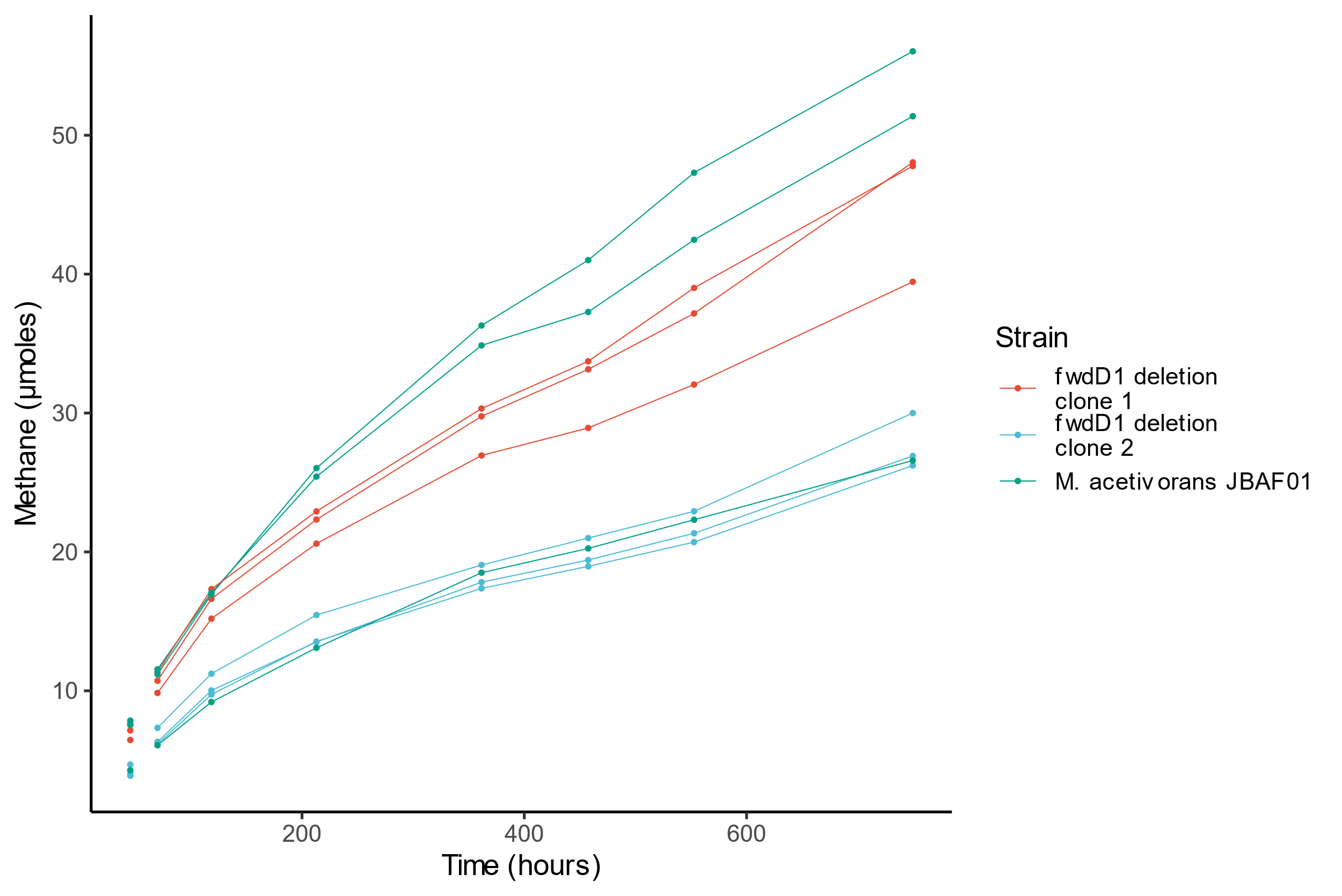

Fig. S4 Methane formation in *M. acetivorans* JBAF01 and *M. acetivorans* JBAF01 ΔfwdD1 on HSF media. Recreation of *fwdD1* deletion had no impact on rate of methanogenesis. Methane measured in three biological replicates. All replicates are shown.

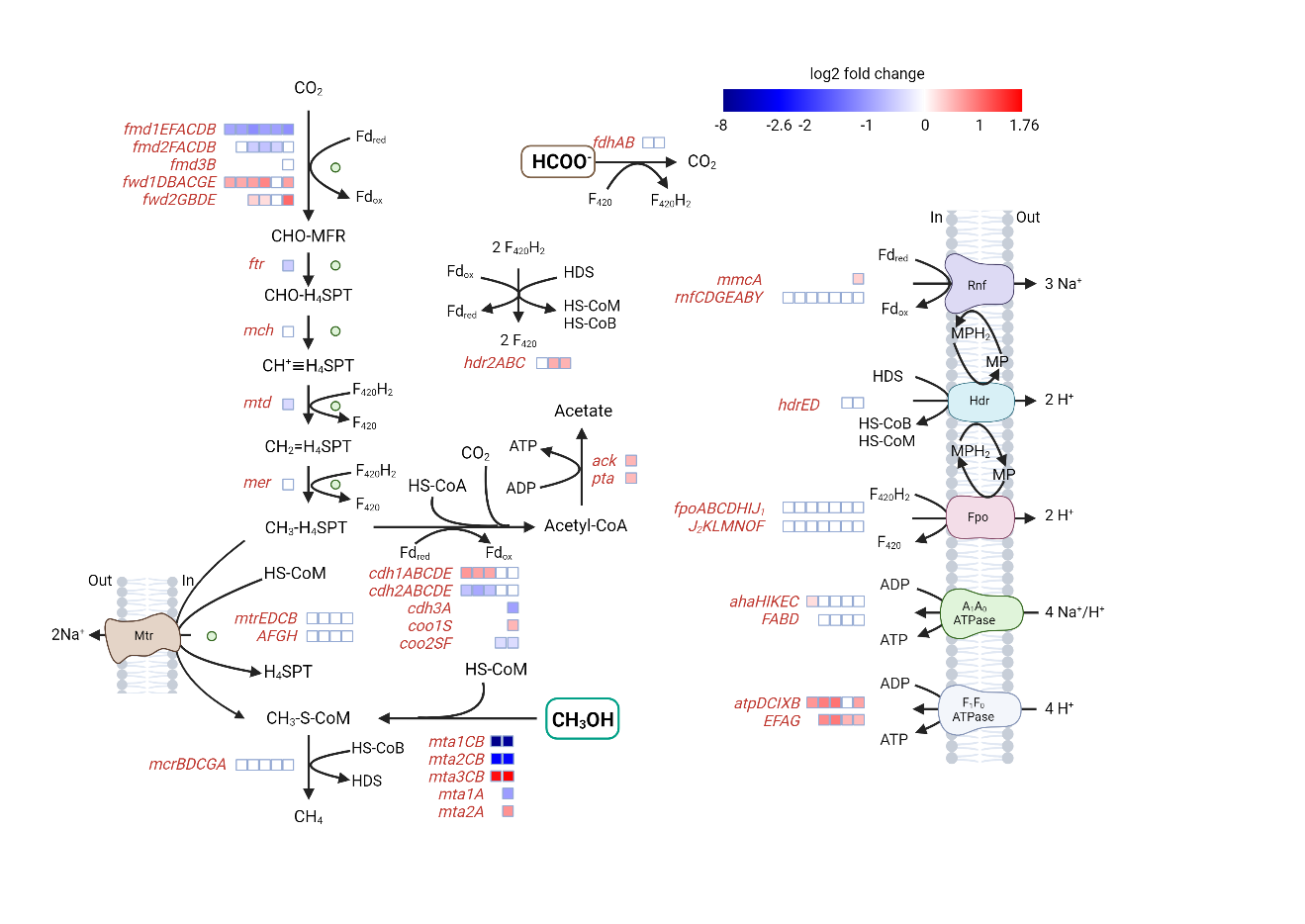

Fig. S5 Methanogenesis pathway in *M. acetivorans* JBAF02 with heatmap showing differential gene expression analysis comparing growth in HSF to growth in HSMe. Log2 fold change indicates the change in the expression level in HSF media compared to HSMe media. The green circle indicates that the reactions are running in reverse when the cell is using methanol as a substrate. Abbreviations: ferredoxin (Fd), methanophenazine (MP), methanofuran (MFR), tetrahydrosarcinapterin (H_4_SPT). Media composition: High salt media with methanol (HSMe), High salt media with formate (HSF). Genes: formylmethanofuran dehydrogenase (*fmd*/*fwd*), formyl transferase (*ftr*), methenyl-H_4_SPT cyclohydrolase (*mch*), methylene-H_4_SPT reductase (*mer*), methyl-H_4_SPT:HS-CoM methyltransferase (*mtr*), methyl-CoM reductase (*mcr*), heterodisulfide reductase (*hdr*), methanol:HS-CoM methyltransferase (*mta*), carbon dioxide dehydrogenase (*cdh*), acetate kinase (*ack*), F_420_H_2_ dehydrogenase (*fpo*). Transcriptomics done on four biological replicates. For genes with multiple homologs, the Gene IDs can be found in SI Appendix text.

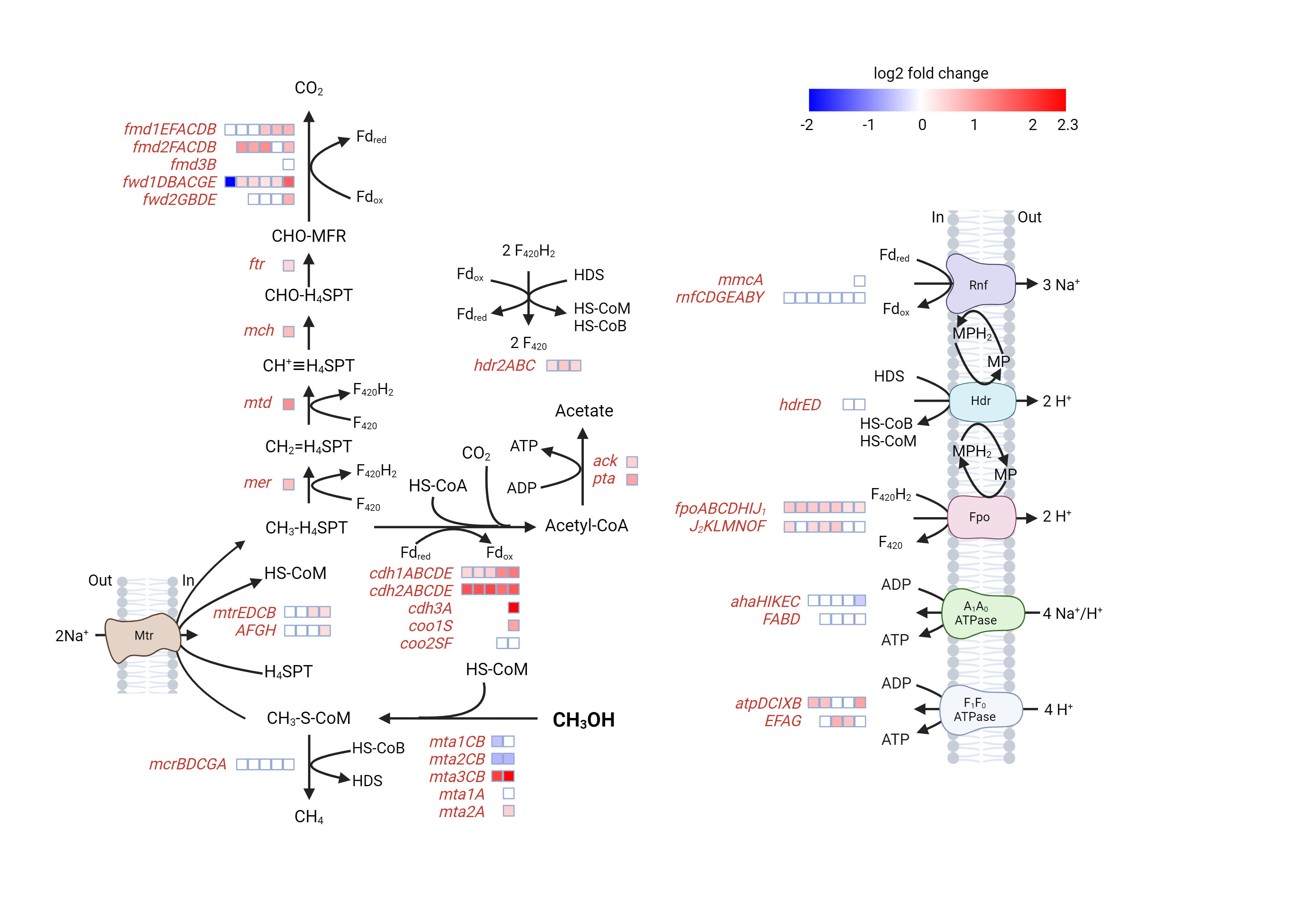

Fig. S6 Differential gene expression analysis comparing *M. acetivorans* JBAF02 growth in HSMe to *M. acetivorans* WWM73 growth in HSMe. Log2 fold change indicates the change in the expression level in *M. acetivorans* JBAF02 compared to *M. acetivorans* WWM73. Abbreviations: ferredoxin (Fd), methanophenazine (MP), methanofuran (MFR), tetrahydrosarcinapterin (H_4_SPT). Media composition: High salt media with methanol (HSMe), High salt media with formate (HSF). Genes: formylmethanofuran dehydrogenase (*fmd*/*fwd*), formyl transferase (*ftr*), methenyl-H_4_SPT cyclohydrolase (*mch*), methylene-H_4_SPT reductase (*mer*), methyl-H_4_SPT:HS-CoM methyltransferase (*mtr*), methyl-CoM reductase (*mcr*), heterodisulfide reductase (*hdr*), methanol:HS-CoM methyltransferase (*mta*), carbon dioxide dehydrogenase (*cdh*), acetate kinase (*ack*), F_420_H_2_ dehydrogenase (*fpo*). Transcriptomics done on three biological replicates for WWM73 and four biological replicates for JBAF02. For genes with multiple homologs, the Gene IDs can be found in SI Appendix text.

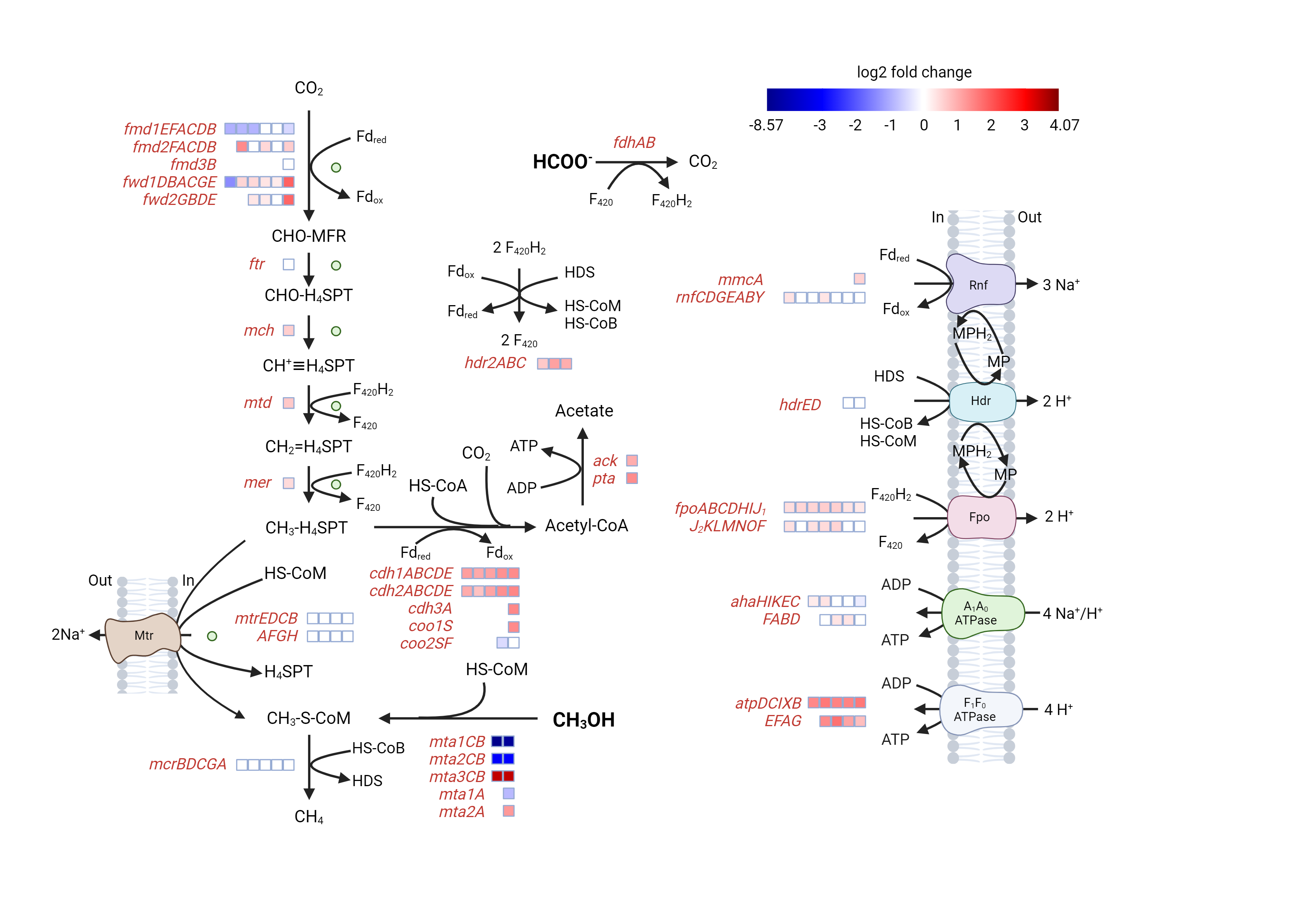

Fig. S7 Differential gene expression analysis comparing *M. acetivorans* JBAF02 growth in HSF to *M. acetivorans* WWM73 growth in HSMe. Log2 fold change indicates the change in the expression level in *M. acetivorans* JBAF02 compared to *M. acetivorans* WWM73. Abbreviations: ferredoxin (Fd), methanophenazine (MP), methanofuran (MFR), tetrahydrosarcinapterin (H_4_SPT). Media composition: High salt media with methanol (HSMe), High salt media with formate (HSF). Genes: formylmethanofuran dehydrogenase (*fmd*/*fwd*), formyl transferase (*ftr*), methenyl-H_4_SPT cyclohydrolase (*mch*), methylene-H_4_SPT reductase (*mer*), methyl-H_4_SPT:HS-CoM methyltransferase (*mtr*), methyl-CoM reductase (*mcr*), heterodisulfide reductase (*hdr*), methanol:HS-CoM methyltransferase (*mta*), carbon dioxide dehydrogenase (*cdh*), acetate kinase (*ack*), phosphate acetyltransferase (*pta*), *Rhodobacter* nitrogen fixation complex (*rnf*), F_420_H_2_ dehydrogenase (*fpo*). Transcriptomics done on three biological replicates for WWM73 and four biological replicates for JBAF02. For genes with multiple homologs, the Gene IDs can be found in SI Appendix text.

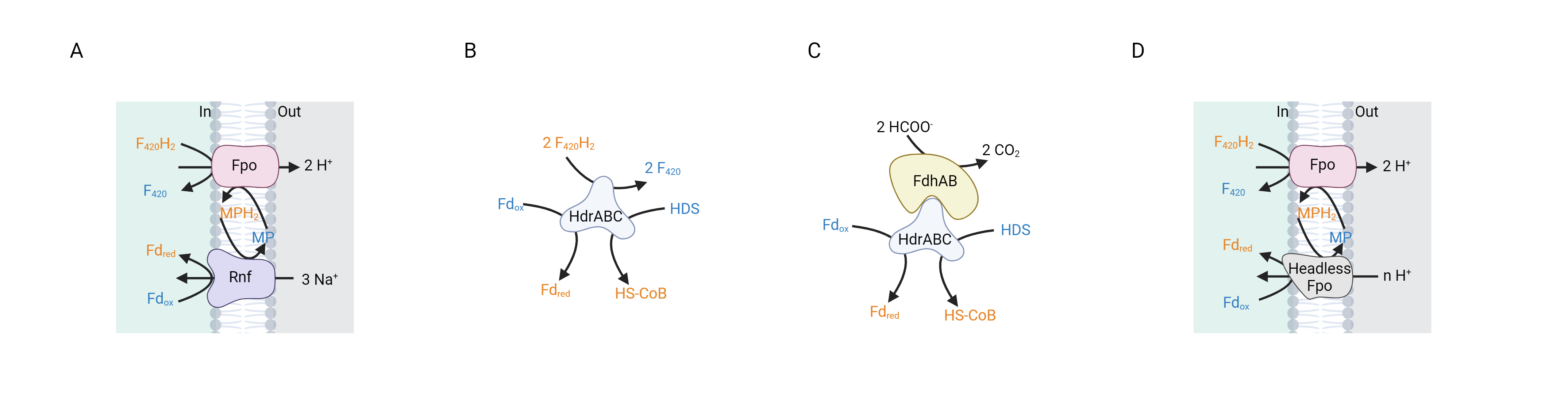

Fig. S8 The four possible pathways for generation of Fd_red_ are presented (A-D) and discussed in detail in SI text. All reduced cofactors are in orange and the oxidized counterparts are in blue. Abbreviations: Coenzyme M (CoM), Coenzyme B (CoB), heterodisulfide between CoM and CoB (HDS), methanophenazine (MP), ferredoxin (Fd), carbon monoxide dehydrogenase (Cdh), methyl-H_4_SPT:coenzyme M methyltransferase (Mtr), F_420_H_2_ dehydrogenase (Fpo), heterodisulfide reductase (Hdr).

Table S1 Formate utilization for methanogenesis in *M. barkeri*.

|  | MF media | | | HSMe media | | |
| --- | --- | --- | --- | --- | --- | --- |
| Time (Hours) | **Methanol (mM)** | **Formate (mM)** | **Methane (µmoles)** | **Methanol (mM)** | **Formate (mM)** | **Methane (µmoles)** |
| 0 | 47.8 | 64.8 | 0.03 | 49.5 | 0.03 | 0.03 |
| 312 | 0 | 38.3 ± 1.2 | 260 ± 16 | 0 | 0.07 | 240 ± 1.5 |

Values at Time = 0 for methanol and formate measured from master mix.
Values at Time = 312 for methanol and formate averaged from three biological replicates
± represents standard deviation

Table S2. Transcriptomics summary

| Condition | Total DEGs | Overexpressed DEGs | Underexpressed DEGs |
| --- | --- | --- | --- |
| *M. acetivorans* WWM73 growth on methanol and formate compared to growth on methanol only | 836/4544 | 376 | 460 |
| *M. acetivorans* JBA01 growth without exogenous acetate and pyruvate compared to growth with exogenous acetate and pyruvate | 1814/4544 | 977 | 837 |
| *M. acetivorans* JBAF02 growth on formate compared to growth on methanol | 1055/4544 | 521 | 534 |

Transcriptomics done on three biological replicates.

Table S3. Biomass yield of *M. acetivorans* on different growth media

| Strain | WWM73 | WWM73 | JBA01 | JBA01 | JBAF02 | JBAF02 |
| --- | --- | --- | --- | --- | --- | --- |
| Medium | HSMe | MF | MFAcP | MF | HSMe | HSF |
| *Y_CH4_* (g total protein mol CH_4_^-1^) | 2.17±0.02 | 2.08±0.03 | 1.67±0.04 | 2.01±0.11 | 1.56±0.10 | 2.48±0.13 |

Values are averaged from three biological replicates.
± represents standard deviation

The total protein was calculated by multiplying OD to the conversion coefficient 69.75 mg L^-^1 OD^-1^. The values were calculated based on the growth in the exponential phase.

Table S4. SNPs found in ALE of *M. acetivorans* JBAF01 to *M. acetivorans* JBAF02

| Position^1^ | Ref. | Change | Present in^2^ | | | | | | | | |
| --- | --- | --- | --- | --- | --- | --- | --- | --- | --- | --- | --- |
| Strain^4^à |  |  | **JBAF01** | **T6** | **T7** | **T18** | **T19** | **T20** | **T28** | **JBAF02^3^** | **Notes** |
| 487691 | T | TC | Y | Y |  |  | Y | Y | Y | Y | Ref (15) |
| 716399 | C | T |  |  |  | Y | Y |  | Y | Y | *ma0610*, archaeal transcription factor B, P285S |
| 941163 | GAAAAA | GAAAAAA | Y | Y |  | Y | Y | Y |  | Y | Ref (15) |
| 1314117 | AGGG | AGG | Y | Y |  |  | Y | Y | Y | Y | Ref (15) |
| 1671119 | ATTTTTTTTT | ATTTTTTTT |  |  |  | Y | Y |  |  | Y | between *ma1403* and *ma1404* |
| 2086881 | T | C | Y | Y | Y | Y | Y | Y | Y | Y | Ref (15) |
| 2086882 | C | T | Y | Y | Y | Y | Y | Y | Y | Y | Ref (15) |
| 2086886 | G | T | Y | Y | Y | Y | Y | Y | Y | Y | Ref (15) |
| 2527716 | A | C | Y |  |  |  |  |  |  |  | *ma2034*, potassium channel protein, N253H |
| 2534542 | GC | GCC | Y | Y | Y | Y | Y | Y | Y | Y | Ref (15) |
| 2836646 | A | G | Y | Y | Y | Y | Y | Y | Y | Y | Ref (15) |
| 2867059 | T | G | Y | Y | Y | Y | Y | Y | Y | Y | Ref (15) |
| 3433198 | ACCC | ACC | Y | Y |  | Y | Y | Y | Y | Y | Ref (15) |
| 3690912 | A | C | Y | Y | Y | Y | Y | Y | Y | Y | *ma2958,* I49M |
| 3746575 | G | A |  | Y | Y | Y | Y | Y | Y | Y | *ma3014*, transcriptional regulator, G22R |
| 4295450 | TCC | TC | Y | Y |  |  | Y | Y | Y | Y | Ref (15) |
| 4638121 | GAAAAAA | GAAAAA |  |  |  |  |  | Y |  |  | *ma3757*, mannosyltransferase B |
| 4664379 | G | A |  |  |  | Y | Y |  | Y | Y | *ma3781*, *rfbM* CDS (mannose-1-phosphate guanylyltransferase (GDP)), S289F |
| 4874563 | CTTTT | CTTT | Y | Y | Y | Y | Y | Y | Y | Y | Ref (15) |
| 4917327 | T | C |  |  |  |  |  |  |  | Y | between *ma4007* and *ma4008* |
| 4945343 | GCC | GCCC | Y |  |  |  |  | Y |  | Y | Ref (15) |
| 5074541 | AGGT | A |  |  |  |  |  | Y |  |  | *ma4158*, *ahaA*, TG67T |
| 5088982 | G | A |  |  |  |  |  | Y |  |  | *ma4167*, D103N |
| 5138462 | T | A |  |  |  |  |  |  | Y |  | *ma4215*, I39F |
| 5601699 | G | T | Y |  |  |  |  |  |  |  | between *ma4550* and *ma4551* |
| 5601701 | C | T | Y |  |  |  |  |  |  |  | between *ma4550* and *ma4551* |

1 : Position in the reference genome (AE010299)
2 : Y indicates Yes, SNP present
3 : Yellow fill indicates that this SNP was not present in the parent strain and was fixed in the population
4 : T indicates the transfer strain as described in (Main text, methods)

Table S5 Growth of *M. acetivorans* WWM73, *M. acetivorans* JBAF02 and *M. maripaludis* J901 in media containing DPI

| Strain | Media | DPI | Growth |
| --- | --- | --- | --- |
| *M. maripaludis* J901 | McFC | No | Yes |
| *M. maripaludis* J901 | McFC | Yes | Yes |
| *M. acetivorans* WWM73 | HSMe | No (Only DSMO) | Yes |
| *M. acetivorans* WWM73 | HSMe | No | Yes |
| *M. acetivorans* WWM73 | HSMe | Yes | No |
| *M. acetivorans* JBAF02 | HSMe | No | Yes |
| *M. acetivorans* JBAF02 | HSMe | Yes | No |
| *M. acetivorans* JBAF02 | HSF | No | Yes |
| *M. acetivorans* JBAF02 | HSF | Yes | No |
| *M. acetivorans* JBAF02 | HSFAcP | No | Yes |
| *M. acetivorans* JBAF02 | HSFAcP | Yes | No |

Experiment done in two biological replicates

Table S6. Ions translocated in methylotrophic methanogenesis compared with the newly engineered metabolisms

|  | Na^+^ translocated CH_4_^-1^ | H^+^ translocated CH_4_^-1^ |
| --- | --- | --- |
| Methylotrophic methanogenesis | 0.33 | 3.33 |
| Formate-dependent methylotrophic methanogenesis | 0 | 4 |
| Formate-dependent CO_2_ reduction methanogenesis^a^ | -1^a^ | 6 |

a : Assumes the hypothesis that Rnf catalyzes reduction of ferredoxin to be true

Table S7. Media composition

| Abbreviation | Methanol [mM] | Formate [mM] | Acetate [mM] | Pyruvate [mM] | Casamino acids [g L^-1^] |
| --- | --- | --- | --- | --- | --- |
| HSMe | 60 | 0 | 0 | 0 | 0 |
| HSF | 0 | 120 | 0 | 0 | 0 |
| MF | 60 | 60 | 0 | 0 | 0 |
| MFAcP | 60 | 60 | 5 | 5 | 0 |
| HSFAcP | 0 | 120 | 5 | 5 | 0 |
| HSFA | 0 | 120 | 5 | 5 | 1 |

Table S8. Strains used in this study

| Name | Description | Reference |
| --- | --- | --- |
| WWM73 | Derived from *M. acetivorans* C2A, Δ*hpt*::P*mcrB-tetR*-ϕC31-int-*attP* | (16) |
| WWM155 | Derived from *M. barkeri* Fusaro, Δ*hpt*::P*mcrB-tetR*-ϕC31-int-*attP* | (16) |
| J901 | Derived from *Methanococcus maripaludis* JJ, Δ*upt* | (17) |
| WWM73-  pJK027A | WWM73 transformed with pJK027A empty vector | This study |
| WWM73-  pJK_Fdh-Mb | WWM73 transformed with pJK_Fdh-Mb plasmid | This study |
| JBA01 | WWM73 ∆*mtrED*(1-44)*::fdh-*T*_fpo_Fusaro_* | This study |
| JBAF01 | WWM73 ∆*frhA*(1001-1371)*DGB*(1-94)*::*P*_mtr_C2A_-fdh-*T*_fpo_Fusaro_* | This study |
| JBAF02 | JBAF01 after adaptive laboratory evolution | This study |
| JBAF01∆fwdD1 | JBAF01 ∆*fwdD1(1-341)* | This study |

Table S9. Primers and gRNA used in this study

| Name | Sequence (5’ -> 3’) |
| --- | --- |
| mtr::fdh P1 | TCCCAGCACAACCCCATGGTC |
| mtr::fdh P2 | GTCGTGGGCACATATTTCAATTCCATTCATTTTCCTCCTTTATAGTTATTAACATGTTAATAAGCTGG |
| mtr::fdh P3 | GAAAATGAATGGAATTGAAATATGTGCCCACGAC |
| mtr::fdh P4 | CCGCAAAATGCAGCTTTACTCAAATTCAATTTTTTTTACAACCTAATTTTTTCCTGTTTTC |
| mtr::fdh P5 | ATTTGAGTAAAGCTGCATTTTGCGG |
| mtr::fdh P6 | GATTTTTTAAGACTCTCTTAAAATTAATCATCCTCTAGTTCTCAGTTTTGGATTCTCAAACCAATAATTC |
| mtr::fdh P7 | CTGAGAACTAGAGGATGATTAATTTTAAGAGAGTCTTAAAAAATC |
| mtr::fdh P8 | ATTAAAAATATATAAAAAAAGGAAACCATATTAGGTTTCCTTTAGTTTTC |
| mtr::fdh P9 | GAAAACTAAAGGAAACCTAATATGGTTTCCTTTTTTTATATATTTTTAATTATAATTGGTGGCGTCCTGATTTCC |
| mtr::fdh P10 | GACCACGTATTCGACGACTTCTGC |
| mtr::fdh P11 | GCAAAAACTGCAGGCGTGGTTG |
| mtr::fdh P12 | GGAAATCAGGACGCCACCAATTATATCTCAGTTTTGGATTCTCAAACCAATAATTC |
| mtr::fdh P13 | TATAATTGGTGGCGTCCTGATTTCC |
| mtr::fdh P14 | CGGCAATAACGACGCTCAATC |
| frh::fdh P15 | GCAATTAACCCTCACTAAAGGGAACAAAAGCTTGACGAAAGTTGTAGAGATCTCTC |
| frh::fdh P16 | CGATGTGCTGGGCGAAGGTG |
| frh::fdh P17 | GCACCTTCGCCCAGCACATCGGCATTAAGCAAGGAGCCCAGATATAAAG |
| frh::fdh P18 same as P2 | GTCGTGGGCACATATTTCAATTCCATTCATTTTCCTCCTTTATAGTTATTAACATGTTAATAAGCTGG |
| frh::fdh P19 same as P3 | GAAAATGAATGGAATTGAAATATGTGCCCACGAC |
| frh::fdh P20 same as P4 | CCGCAAAATGCAGCTTTACTCAAATTCAATTTTTTTTACAACCTAATTTTTTCCTGTTTTC |
| frh::fdh P21 same as P5 | ATTTGAGTAAAGCTGCATTTTGCGG |
| frh::fdh P22 | TCTCAGTTTTGGATTCTCAAACCAATAATTC |
| frh::fdh P23 | GAATTATTGGTTTGAGAATCCAAAACTGAGATTGCAGGCAGGTACTTAAAGCC |
| frh::fdh P24 | CTATTTTCCGAAGATTTTTTAAGACTCTCTTAAAATTAATCATCCTCTAGTTGCCGCCGTCCTGTGCCTTCT |
| frh::fdh P25 | GCAACTAGAGGATGATTAATTTTAAGAGAGTCTTAAAAAATC |
| frh::fdh P26 same as P8 | ATTAAAAATATATAAAAAAAGGAAACCATATTAGGTTTCCTTTAGTTTTC |
| frh::fdh P27 | GGAAACCTAATATGGTTTCCTTTTTTTATATATTTTTAATAGCTCCTCCAGTGGTGAAGG |
| frh::fdh P28 | AGGGCGAATTGGGTACCGGGCCCGGGCCTTCTGTTCCTCATTC |
| 0AB P29 | CGAAGCTGCTGGTGAAAGAGAACCTATCTTACCTGCTAAAATCTAAG |
| 0AB P30 | CTGGAGCCGGTGAGCGTGGTTCTCGCGGTATCATTGCAGC |
| 0AB P31 | GAGAACCACGCTCACCGGCTCCAGATTTATC |
| 0AB P32 | ACTGCTGATACGTTGAGGTCACCCCATGTACTTCTTTTATTGTACTC |
| 0AB P33 | GTACATGGGGTGACCTCAACGTATCAGCAGTATGC |
| 0AB P34 | GAGAATCACTCCTATTTTTTTGATATATACATCATAAC |
| 0AB P35 | GTTATGATGTATATATCAAAAAAATAGGAGTGATTCTCATGACTGAATATAAACCTACCGTTAGG |
| 0AB P36 | CGGGCGCTGCGGGTCGTGGGGCGGGCGTTATGCTCCAGGTTTTCTTGTCATG |
| 0AB P37 | GCCCCACGACCCGCAGCGCCCG |
| 0AB P38 | GTAAGATAGGTTCTCTTTCACCAGCAGCTTCG |
| fwdD gRNA P39 | GAGGGTCTCAAAGTGTACTGGACCCTGACACCCAGTTTGGAGACCCAC |
| fwdD gRNA P40 | GTGGGTCTCCAAACTGGGTGTCAGGGTCCAGTACACTTTGAGACCCTC |
| fwdD P41 | TGTCACCTGCAGCAGCCGCATTATAGTGCCCGCGCATTC |
| fwdD P42 | CAGATGAGGAATTTCCCCTGGAAATGAAGGCCCTGATG |
| fwdD P43 | CATTTCCAGGGGAAATTCCTCATCTGGGGTGTAAC |
| fwdD P44 | GCAACACCTGCTTCAGGGAGAAAAAATGATTTTAGCATGAAACACAGC |
| GGA P61 | GTAATTATAACCCGGGCCCTATATATGGATCCCGCATATGTTTTTTCTAAACTCTCATTTG |
| GGA P62 | CCTGAGCCAGCTTGGAGGAGATTAAGAACTAATTTTTAAATTAATTCTTTTTTGATACTGC |
| GGA P63 | ATCTCCTCCAAGCTGGCTCAGG |
| GGA P64 | CCCAAAATGATTTTAATAAATTAAGGAGGAAATTCATATGGATAAGAAGTACTCAATTGGACTTG |
| GGA P65 | ATGAATTTCCTCCTTAATTTATTAAAATCATTTTGGG |
| GGA P66 | GAAATGTATTTATTCAAGTACTGCATCTAGGCGCCTGATGCGGTATTTTCTC |
| GGA P67 | CCTAGATGCAGTACTTGAATAAATACATTTC |
| GGA P68 | CGAGACCGTGCGGCCGCTAGGTCTCTACTTCTAACTCCCCTTAATTTAATTGGAATTC |
| GGA P69 | AGAGACCTAGCGGCCGCACGGTCTCGGTTTTAGAGCTAGAAATAGCAAGTTAAAATAAG |
| GGA P70 | AAAAGCACCGACTCGGTGCCAC |
| GGA P71 | GTGGCACCGAGTCGGTGCTTTTGCCGAACCCTCCGGTTCTC |
| GGA P72 | CGGTACCTCTAGAACTATAGCTAGCATAACTTCGTATAATGTATGCTATACGAAGTTATGCCTTTTAAAAAGGGATTGAGCG |
| GGA P73 | CATACATTATACGAAGTTATGCTAGATGCGCAGGTGAGATCCGCACCTGCATTGACCGGTTGTTAACGTTAGCCGGCTAC |
| GGA P74 | GCTAACGTTAACAACCGGTCAATGCAGGTGCGGATCTCACCTGCGCATCTAGCATAACTTCGTATAATGTATGCTATACG |
| GGA P75 | GGTAATTATAACCCGGGCCGATGCGCAGGTGAGGATCCGCACCTGCATTGTCCCGCATATGTTTTTTCTAAACTCTC |
| GGA P76 | GAGAGTTTAGAAAAAACATATGCGGGACAATGCAGGTGCGGATCCTCACCTGCGCATCGGCCCGGGTTATAATTACC |
| gRNA | |
| Target gene | **gRNA (5' -> 3')** |
| *fwdD1* | GTACTGGACCCTGACACCCA |

Table S10. Plasmids used in this study

| Name | Description | Reference |
| --- | --- | --- |
| pM000 | *E.coli*/*M. acetivorans* shuttle vector, containing *pac* for selection on puromycin media | (18) |
| pM001 | Derived from pM000 containing *hpt* for counterselection on 8-aza-2,6-diaminopurine media | (18) |
| p425GPD | Cloning vector, used in pfdh V3 construction | (19) |
| pNB730 | Integrative vector for WWM73, used as PCR template | (20) |
| pJK027A | Integrative vector for WWM73, used for formate dehydrogenase (Fdh) in vitro test | (16) |
| pMSJB000AB | Derived from pM000, BsaI stie-free, codon optimized *pac* | This study |
| pMSJB001AB | Derived from pM001 BsaI site-free, codon optimized *pac* | This study |
| pGGA2 | Derived from pMSJB001AB, that contains the Cas9 and gRNA expression cassette, it features *Bsa*I sites for gRNA integration and *PaqC*I sites for repair fragment insertion. | This study |
| pJK_Fdh-Mb | Derived from pJK027A, the gene of Fdh is under the control of P*_mcrB(tetO1_*_)_ | This study |
| pfdh V3 | Carrying the integration fragment containg an upstream homologous recombination arm (*frhA* (1-1000)), P*_mtr_C2A_*-*fdh*-T*_fpo_Fusaro_*, a 400 bp DNA fragment (*frhG* 494 - *frhB* 94) for the 2nd round marker removal, the *pac-hpt* marker cassette, a downstream homologous recombination arm (*frhA* 1355 - *frhG* 493) | This study |
| pGGA2-fwdD1 | Derived from pGGA2 for *fwdD1* deletion | This study |

Dataset S1 (separate file). Differential gene expression analysis of *M. acetivorans* WWM73 growing on HSMe compared to MF

Dataset S2 (separate file). Differential gene expression analysis of *M. acetivorans* JBA01 growing on MFAcP compared to MF

Dataset S3 (separate file). Differential gene expression analysis of *M. acetivorans* JBAF02 growing on HSMe compared to HSF

Dataset S4 (separate file). Differential gene expression analysis of *M. acetivorans* WWM73 growing on HSMe compared to *M. acetivorans* JBAF02 growing on HSMe

Dataset S5 (separate file). Differential gene expression analysis of *M. acetivorans* WWM73 growing on HSMe compared to *M. acetivorans* JBAF02 growing on HSF
